# Rapid and reliable quantification of cytosolic mRNA escape (RNASCAPE)

**DOI:** 10.64898/2026.04.07.716953

**Authors:** Frank H. Schulz, Emily Winther Sørensen, Steen W.B. Bender, Artu Breuer, Georgios Kyriakakis, Marcus W. Dreisler, Georgios Bolis, Athanasios Oikonomou, Konstantinos Tsolakidis, Stefanos Arampatzis, Guangjun Nie, Nikos S. Hatzakis

## Abstract

Endosomal escape of mRNA remains a critical bottleneck in oligonucleotide therapeutics, with typically less than 5% of delivered mRNA reaching the cytosol. Precise quantification of this escape is hindered by stochastic variability in uptake, release, and expression, while existing methods lack scalability and accuracy. Here we introduce RNASCAPE, a deep learning framework trained on biologically realistic simulations that estimates unlabelled mRNA cytosolic escape efficiency using only three timepoints of EGFP reporter expression and four lipid nanoparticle (LNP) meta-parameters. Benchmarking on synthetic data shows RNASCAPE achieves a mean absolute percentage accuracy of 78%. RNASCAPE accurately predicts escape efficiencies around 5–9% across diverse cell types and revealed that exchanging cholesterol to β-sitosterol drastically reduces mRNA loading and enhances twofold its functional release. By enabling robust, microscopy-agnostic quantification without requiring specialized cell lines or labeled cargo, RNASCAPE provides a scalable framework for benchmarking and rational design of LNP formulations, advancing nucleic acid therapeutic delivery.

## INTRODUCTION

The limited endosomal escape of oligonucleotide therapeutics in cytosol remains a bottleneck restricting their widespread implementation. Lipid nanoparticles (LNPs) have emerged as a leading platform enabling the success of mRNA vaccines and siRNA and other nucleotide-based pharmaceutics owing to their ability to encapsulate and protect nucleic acids while enabling cellular uptake. Their ionizable lipids are designed to promote LNP fusion with the maturing endosome and release the cargo into the cytosol. In practice, however, only a small fraction of internalized mRNA or siRNA molecules escape from endosomes, typically estimated to be from ∼5% (1–4) to ∼15% (5,6) of the delivered material.

Endosomal escape is inherently stochastic, transient, and both spatially and temporally heterogeneous within cells as it is dependent on LNP properties and an interplay of several biological processes (1). LNPs can enter cells via multiple endocytic routes that can vary dependent on cell type, their composition, and protein corona effects (7–11). Once internalized, LNPs may release cargo during endosome maturation, undergo lysosomal degradation, or recycle to the plasma membrane (Fig. 1a top). For cytosolically released mRNAs, translation depends on mRNA stability, translation efficiency, and protein turnover, all of which vary with cellular state resulting in high inter-cell variation (Fig. 1a bottom). The co-existence of these processes, each with distinct rate constants, challenges disentangling their effects analytically and current state-of-the-art methods primarily focus and address only parts of the problem (12).

**Fig. 1:**
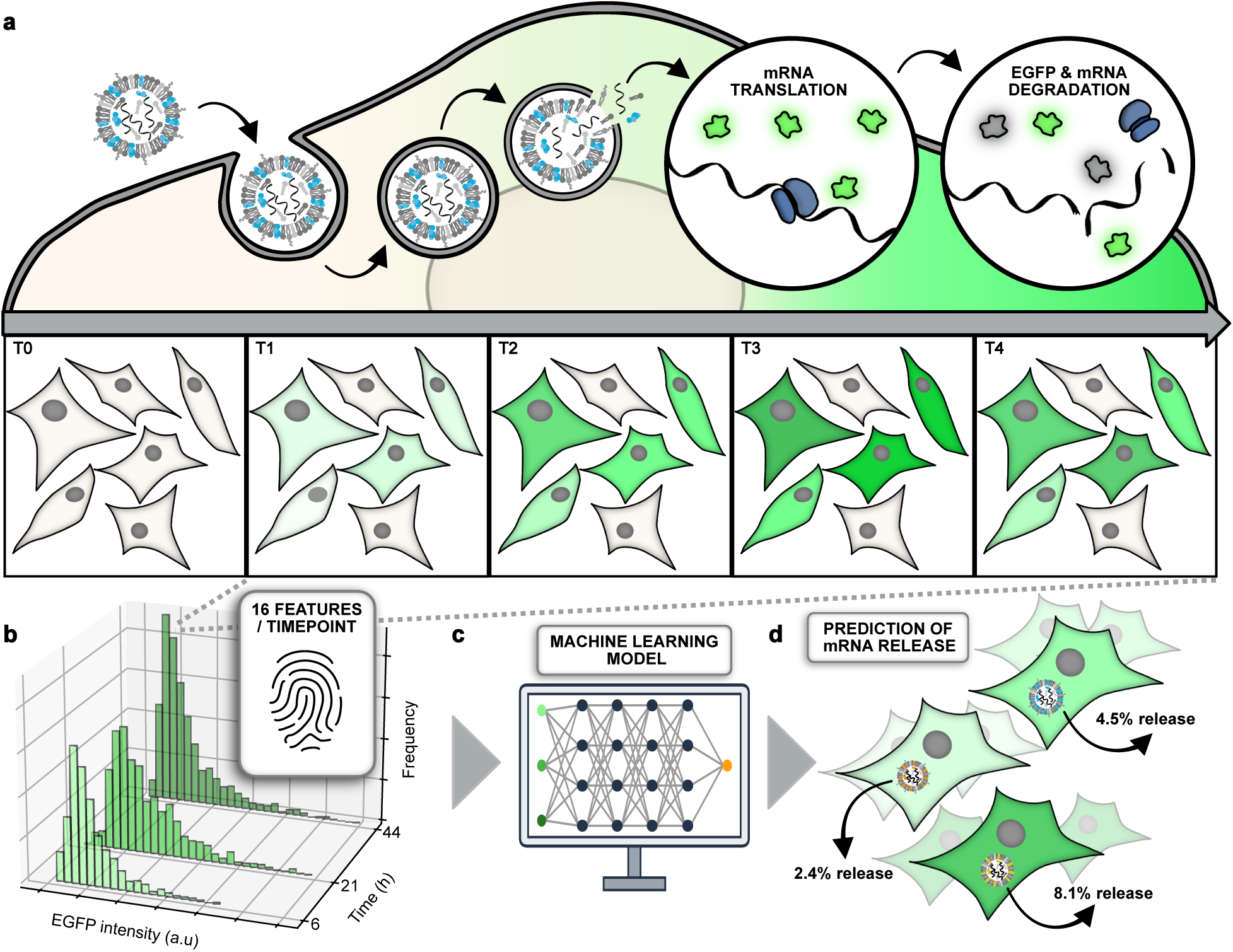
Cartoon representation of deep learning assisted quantification of released mRNA from internalized LNPs relying exclusively on 3 timepoint readouts and LNP characterization. **a** top, Schematic representation of cellular uptake of LNPs that upon mRNA escape into cytosol lead to expression of EGFP cells. a bottom, Representation of time dependent EGFP expression upon LNP addition: LNPs enter cell through endocytosis (T0). As an LNP-containing endosome matures, endosomal membrane is expected to be disrupted by LNP partially releasing mRNA. Escaped mRNA is then continuously translated to EGFP molecules, increasing the cell‘s fluorescence intensity (T1-T3). Both mRNA and EGFP are eventually degraded, resulting in a peak in fluorescence intensity (T3), followed by a decrease in fluorescence intensity (T4). **b,** Quantification of expression levels of EGFP in single cells using fluorescence-based microscopy techniques. Each distribution is derived from single cell recordings at 3 distinct, but flexibly separated within 48h, timepoints. Each EGFP expression distribution has its own unique distributional fingerprint that is described by 16 features per timepoint that are used for predicting the mRNA release ratio. **c,** Deep learning model trained on synthetic simulated data accounting for wide distribution of relevant parameters. The trained model takes as input the experimentally derived EGFP expression at three flexibly separated timepoints in the span of two days along with four experimentally descriptive meta-parameters, LNP dimension, loading efficiency, number of internalized LNPs, and GFP degradation rate. **d,** The model predicts the mRNA release ratio of individual LNP compositions.

Bulk reporter protein expression assays, while scalable, convolve multiple interdependent factors including LNP uptake, size heterogeneity, fraction of loaded and loading efficiency, the probability of endosomal escape, mRNA stability, translation efficiency, and cell division (13,14). Single-particle imaging methods have been developed to probe escape directly, yet they have not realized their full potential due to significant limitations. Electron microscopy provides high spatial resolution and can visualize pore formation and rare cytosolic release events (1) but requires fixation, uses non-native probes like gold-nanoparticles, and has inherently low throughput. Fluorescent galectin-9 recruitment offers a live-cell compatible and high throughput assay of endosomal rupture and can be coupled to reporter expression (15). It however requires specialized cell lines (4), limiting its generic applications across new cells, only reports gross endosomal damage and provides a qualitative measure of escaped mRNA molecules. Single-molecule fluorescence in situ hybridization (smFISH) and super resolution microscopy can directly visualize cytosolic mRNA molecules and offer nanoscale insight into escape sites such as endosomal tubules, respectively (5,6,13). However, they are fixation-based, laborious, and incompatible with functional readouts and live cell dynamics. Split luciferase complementation can quantify endosomal release of pH-responsive polymer nanoparticles, albeit it requires specialized cell lines and is not currently compatible with mRNA–LNP systems (16).

Modelling and simulations have provided mechanistic insight on intracellular transport and release, but their quantitation remain limited. Stochastic models (17), and differential equation-based mathematical frameworks (18–20) elucidate rate-limiting steps but rely on bulk constraints and do not provide single-cell escape estimates. Multiparameter models of siRNA delivery have similarly quantified escape (21) but bypass the added layers of translation and protein stability inherent to mRNA. Modelling of time-resolved single-cell microscopy has extracted onset-time distributions and expression kinetics (22), offering detailed temporal insight, but even these models do not yield quantitative estimates of cytosolic release ratios. Kinetic approaches have also been applied at the formulation level, where a simple ODE-based model connected ionizable lipid chemistry to population-level siRNA efficacy (23), but this remains bulk and cannot resolve single-cell heterogeneity. More recently, probabilistic frameworks have been used to estimate siRNA escape from live-cell imaging (24), but these rely on labelled cargo, are tailored to siRNA, and are not generalizable to mRNA–LNP systems. Machine learning applied to cell imaging can identify productive entry routes and guide rational LNP design (25), but such approaches still report functional output rather than quantifying how large a fraction of mRNA molecules reach the cytosol. Collectively, these efforts have advanced mechanistic understanding, but none disentangle the coupled stochastic processes of uptake, escape, and expression from minimal single-cell input data.

Here, we provide RNASCAPE, a platform that estimates intracellular endosomal escape of LNP-delivered mRNA from live-cell microscopy data. The model disentangles the contributions of LNP heterogeneity, uptake, endosomal escape, and downstream expression, and accurately quantifies endosomal escape of LNP-delivered mRNA to be approximately 5–9% depending on LNP sterol content. Implementation of RNASCAPE across formulations and HEK293 and HeLa cells revealed that replacing cholesterol with β-sitosterol radically reduced mRNA loading in LNPs and enhanced functional mRNA delivery two-fold. RNASCAPE operates across diverse microscopy setups and cell lines and was validated using both spinning disc confocal and lattice light-sheet datasets, demonstrating its robustness and adaptability. Importantly It does not require specialized cell lines or labelled mRNA and only needs three experimental readouts and four meta-parameters to output precise endosomal escape estimates. To ensure ease of use for a broad audience, we provide a standalone executable version of RNASCAPE, which requires only a set of time-dependent features and meta-parameters to automatically and rapidly quantify endosomal escape. RNASCAPE establishes a standardized framework for benchmarking and high-throughput screening of LNP formulations, enabling rational design and comparative studies to advance oligonucleotide delivery platforms.

## RESULTS

RNASCAPE is a framework tool that estimates mRNA cytosolic escape efficiency using only three flexibly separated within 48h timepoints of EGFP reporter expression and four experimentally acquired LNP meta-parameters (Fig. 1).

### Simulation and model

To address the challenge of underdetermined parameter inference from fluorescence-based measurements, we generated biologically grounded simulations that reflect biological realism spanning plausible parameter ranges (20) and trained a model on the high-dimensional structure of the data (Supplementary Fig. 1). The simulator produced ∼800,000 experiments that each contain 10–1000 single-cell EGFP expression time courses by sampling 12 kinetic parameters that capture delivery and expression: mRNA release percentage, biological noise, cell size, expression rate, cell doubling rate, mRNA half-life, EGFP half-life, measurement time, number of cells, internalized LNPs per cell, mRNAs per LNP, and the percentage of empty LNPs (Fig. 2a, Methods and Tables 1–2) (20,22). Each kinetic parameter was drawn from a biologically relevant uniform range based on literature assumptions (see Methods and Tables 1–2) (18,26). The simulator resolves dynamics at the level of single cells, individual mRNAs, and individual EGFP molecules, with values independently sampled to produce distributional variability that mirrors biological experiments (see Methods).

**Fig. 2:**
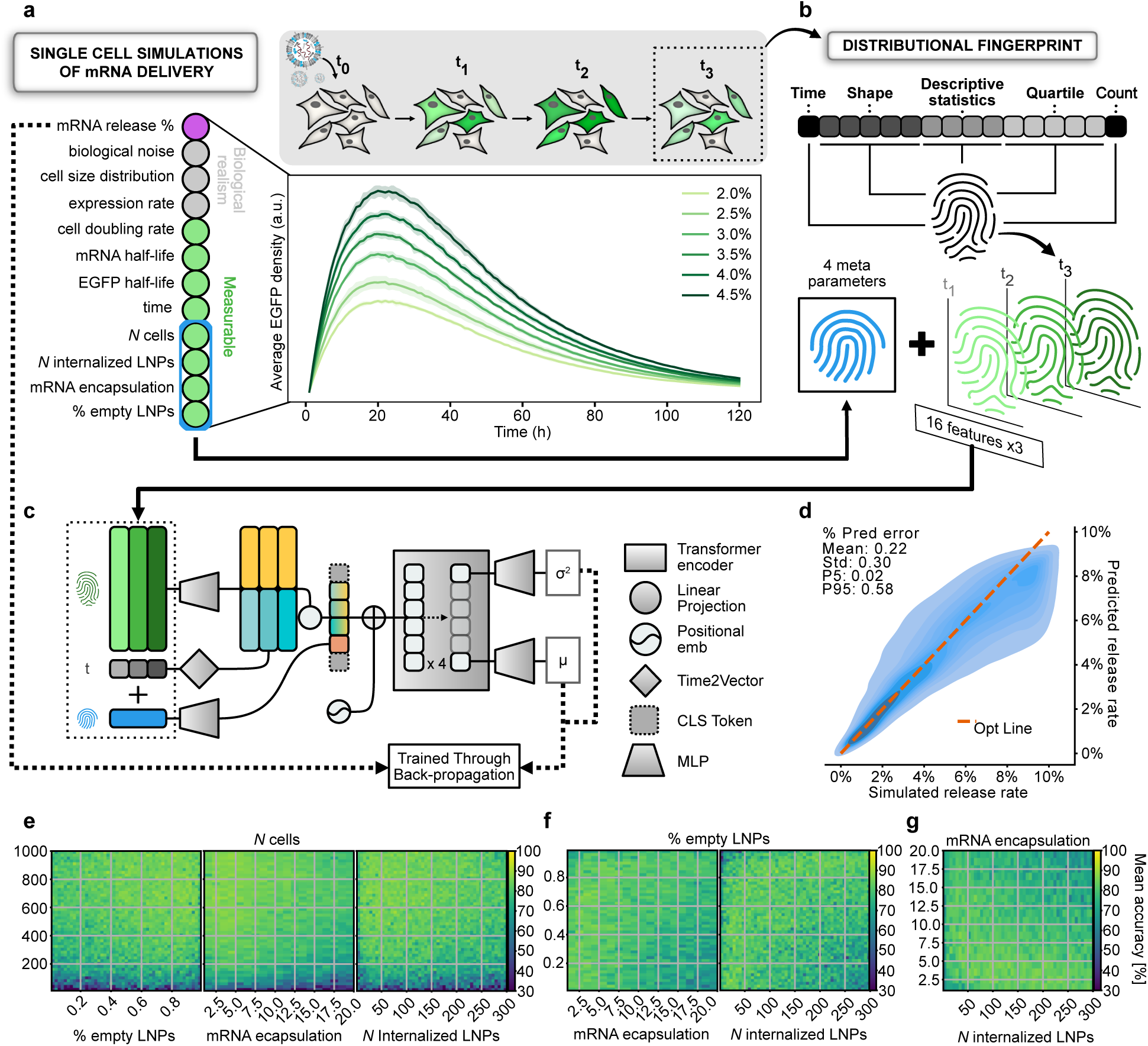
Training, architecture, and evaluation of deep learning model performance. **a,** Training data generation. Single-cell EGFP time courses are simulated by sampling 12 kinetic parameters that capture biological realism. Replicates of single-cell datasets are generated for each defined release ratio to provide training distributions for the model. Plots show the mean EGFP per cell with std across three simulated experiments, with an initial rise followed by a decay driven by mRNA and EGFP degradation. **b,** Feature extraction. At each timepoint the single-cell expression distribution is reduced to 16 features, termed a fingerprint of cell expression distribution which is used for inference. Features encode the acquisition time and cell count as well as distribution shape, including location and scale statistics, quantiles, skewness, and tail weight. The three fingerprints together summarize how the population evolves over time. **c,** Model architecture. The model takes as input four meta-parameters together with expression distributions from three timepoints. Inputs are first processed separately (meta-parameters, time of measurement, and expression distributions). Features and meta-parameters are enriched through individual MLPs, while a temporal encoding is extracted from the timepoints using a small Time2Vec model. The temporal encoding is added to the enriched features through a linear projection before concatenated with the enriched meta-parameters and two classification (CLS) tokens, forming a six-entry data sequence. Positional encoding is added before passing the sequence through a transformer encoder. The CLS tokens are then processed by separate MLPs to predict the mean release ratio and log-variance. During training, simulated targets serve as ground truth, enabling optimization via backpropagation with a heteroscedastic loss. **d,** Model performance on an isolated test dataset. Predicted versus target release ratios are displayed as a kernel density plot with the diagonal marking perfect agreement. **e–g,** Heatmaps displaying mean model accuracy (1 - relative error) dependance on experimental meta-parameters. Data computed on the held-out test across pairs of meta-parameters. All heatmaps are normalized and share a global pseudocolor bar starting from the global minimum value. **e,** Model accuracy dependance on % of empty LNPs, mRNA encapsulation and *N* of internalized LNPs. All heatmaps display *N* cells on the y-axis versus each of the other meta-parameters on the x-axis. **f,** Heatmaps with % empty LNPs on the y-axis versus mRNA encapsulation (left) and *N* of internalized LNPs respectively on the x-axis. **g,** Heatmap of mRNA encapsulation (mRNAs per loaded LNP) on the y-axis versus *N* internalized LNPs per cell on the x-axis.

**Table 1:** Feature extractions and explanations for each. Each of these features is extracted for each measuring time, accumulating to 48 features for our presented 3 timepoint measures.

| Feature | Explanation |
| --- | --- |
| Count | Amount of units (cells) present in the measure |
| Mean | Average expression level<br>$\mu = \sum_{i=1}^N x_i / N$ |
| Standard Deviation | Expression spread around the mean (assuming Gaussian statistics)<br>$\sigma = a \sqrt{\frac{\sum_{i=1}^N (x_i - \mu)^2}{N}}$ |
| Min value | Minimum present expression value |
| Max value | Max present expression value |
| Percentile (10%, 25%, 50%, 75%, 90% ) | Expression value at the given percentile through the sorted expression (50% à Median) |
| Skewness | Expression distribution unbiased asymmetric behavior. |
| | $skew a = \frac{\sqrt{N(N-1)}}{N-2} \frac{m_3}{m_2^{3/2}}$ $m_i = \frac{1}{N} \sum_{n=1}^N (x_n - \mu)^n$ |
| Kurtosis | Explains the unbiased widening of the expression distribution.<br>$kurtosis = \frac{n(n+1)}{(n-1)(n-2)(n-3)} * \frac{\sum (x_i - \bar{x})^4}{s^4} - \frac{3(n-1)^2}{(n-2)(n-3)}$ |
| Boxcox Lambda | Finds $\lambda$ that maximizes the log-likelihood of the Box-Cox-transformed data being normally distributed.<br>Boxcox transformation:<br>$y_i(\lambda) = \begin{cases} \frac{x_i^\lambda - 1}{\lambda}, & \text{if } \lambda \neq 0 \\ \ln(x_i), & \text{if } \lambda = 0 \end{cases}$ |
| Log Mean | Log transforms the data before extracting the mean |
| Log Standard Deviation | Log transforms the data before extracting the standard deviation |
| Measuring Time | Time of measure in h since start |

**Table 2:**
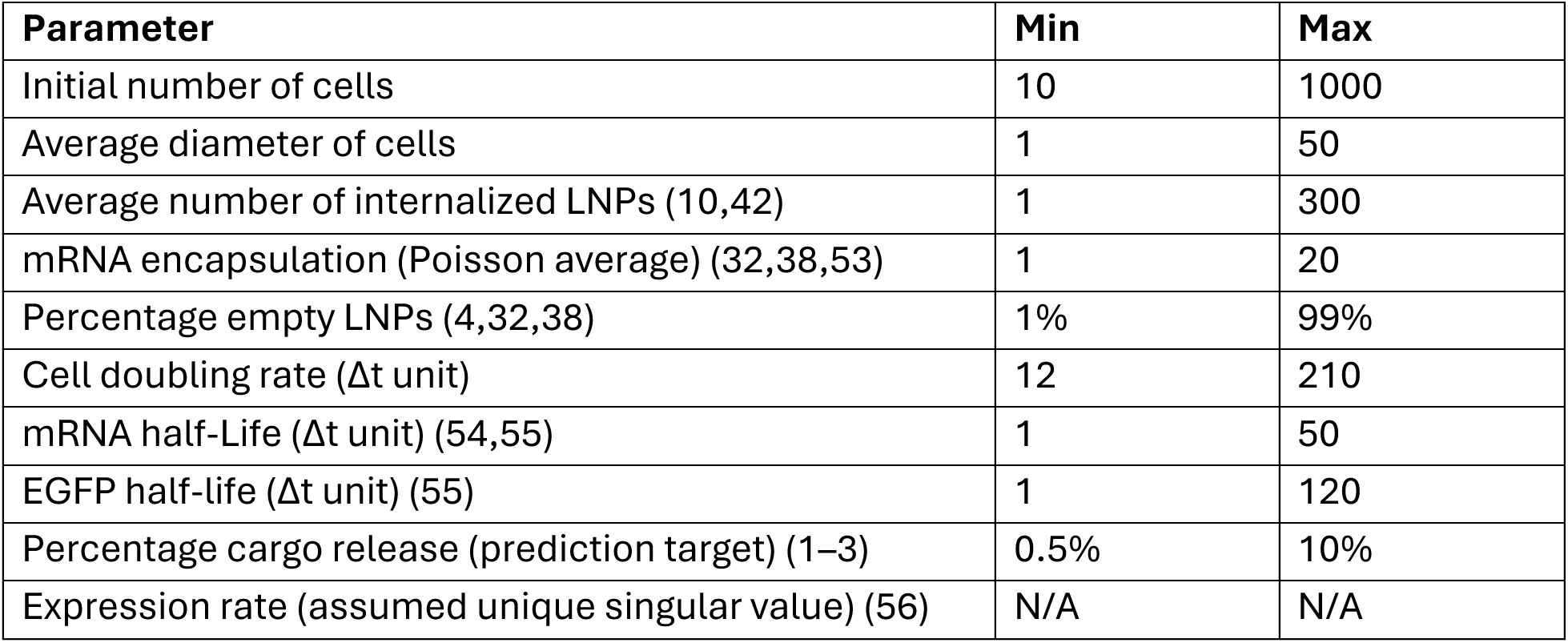
Overview of parameter ranges used in data simulations schematics. . Each parameter range is uniformly distributed and supported through literature.

| Parameter | Min | Max |
| --- | --- | --- |
| Initial number of cells | 10 | 1000 |
| Average diameter of cells | 1 | 50 |
| Average number of internalized LNPs (10,42) | 1 | 300 |
| mRNA encapsulation (Poisson average) (32,38,53) | 1 | 20 |
| Percentage empty LNPs (4,32,38) | 1% | 99% |
| Cell doubling rate ($\Delta t$ unit) | 12 | 210 |
| mRNA half-Life ($\Delta t$ unit) (54,55) | 1 | 50 |
| EGFP half-life ($\Delta t$ unit) (55) | 1 | 120 |
| Percentage cargo release (prediction target) (1–3) | 0.5% | 10% |
| Expression rate (assumed unique singular value) (56) | N/A | N/A |

To capture distributional variability and stochastic noise at each timepoint, the single-cell expression distribution is compressed to a 16-feature “expression fingerprints” capturing time, cell count, mean, variance, quantiles, and shape (skewness/kurtosis) (Fig. 2b) that is used for inference. This encoding preserves both population-level variability and rare outlier behaviors, providing the model with a rich statistical summary of each experimental state.

The model predicts mRNA endosomal escape from three experimental expression distributions and four measured meta-parameters: average mRNAs per loaded LNP, fraction of empty LNPs, internalized LNPs per cell, and number of cells at start of experiment (Fig. 2c). Training randomized sampling intervals and ensured that any three timepoints within 48 h, provided that the first timepoint occurs with the first 6 h, are valid.

Inputs are organized into meta-parameters, measurement times, and fingerprints, enabling the model to disentangle uptake, escape, and expression and to return standardized estimates of cytosolic release. Meta-parameters and fingerprints are independently enriched through separate multilayer perceptron (MLP), with each fingerprint processed individually to project into a shared representation space. To capture temporal dependencies, the three measurement times are encoded via a Time2Vec (27) module and projected into their corresponding enriched fingerprints. The resulting temporal-enriched features are concatenated with the encoded meta-parameters and two randomly initialized classification (CLS) tokens. Each timepoint, together with the meta-parameters and CLS tokens, forms a seven-token sequence, which is processed by a four-layer transformer encoder. Given the inherent biological variability and noise in expression data, the architecture leverages global multi-head attention, enabling adaptive weighting of inputs based on context. Finally, the two CLS tokens are passed to separate MLPs that output the predicted release ratio and log-variance.

### Model validation on ground truth synthetic data

To assess the model’s performance and stability, we evaluated RNASCAPE on an intendent test set comprising 10% of the simulated data withheld during training. Predicted versus target release ratios aligned closely along the ground truth identity line with an average relative error of 0.22 ± 0.30. Predictions were most tightly clustered at lower release ratios (<4%) and became more dispersed at higher values (Fig. 2d). Because experimental meta-parameters must often be estimated, we further examined model sensitivity to parameter misspecification. Perturbation analysis revealed that overestimating the empty-LNP fraction and underestimating initial cell numbers exerted the greatest influence on predictive accuracy (Supplementary Figs. 2–7), whereas other parameters had minor effects.

We next quantified how measurable inputs influence model accuracy by calculating the mean accuracy (1 − relative error) across all pairwise combinations of meta-parameters in the test set (Fig. 2e–g). As expected, accuracy was low for low cell counts (<60% at <200 cells), at high loading rates (<60% at >20 mRNA per LNPs and at extreme high fraction of empty LNPs (<60% at >90% empty LNPs). Across biologically relevant values of meta and experimentally determined values of empty-LNP fraction, mRNA cargo loading, accuracy remained high, with a general peak at ∼85%, demonstrating the model’s robustness.

### Quantification of mRNA escape on experimental data

Application and experimental validation of the model operates in three main steps (Fig. 3a). Step 1 quantifies the fraction of empty LNPs and the mean number of mRNA molecules per LNP by single-particle imaging. Step 2 determines the average number of LNPs internalized per cell under identical conditions to those used for EGFP expression. Step 3 quantifies single cell EGFP expression levels at three distinct time points following LNP transfection.

**Fig. 3:**
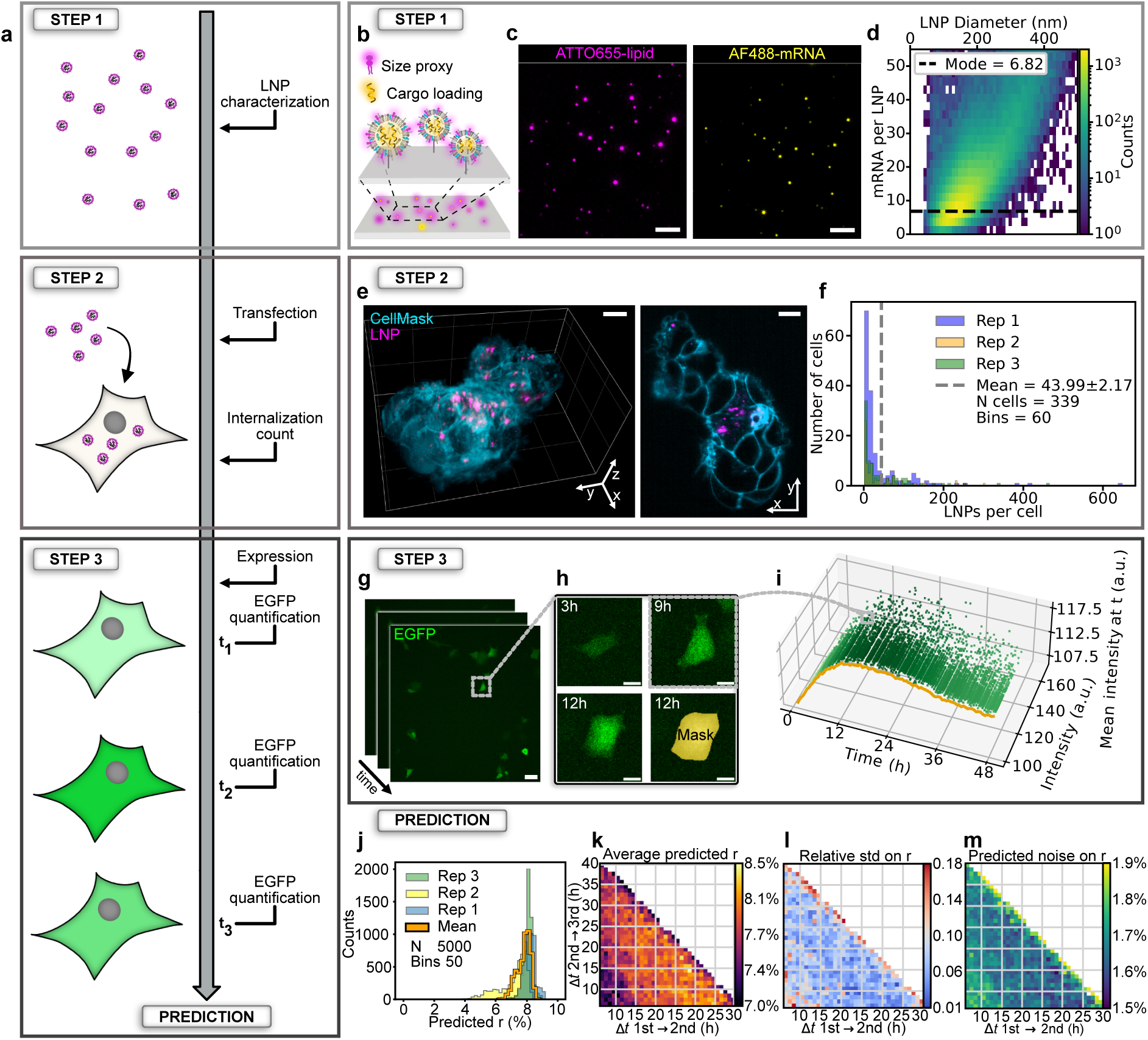
Model and method validation on spinning disc experimental data. **a**, mRNA endosomal escape quantification involves three experimental steps: (1) single-particle characterization of LNP’s heterogeneity and loading, (2) determination of internalized LNPs per cell, and (3) time-resolved EGFP expression in transfected cells. Step 1 (**a**–**c**). **b**, Single-particle characterization determines LNP size distribution, mRNA copy number per LNP, and the ratio of loaded to empty particles. LNPs are labeled with fluorescently labelled lipids (magenta) and mRNA (yellow) and tethered on passivated glass surface. **c**, Typical TIRF micrographs displaying the lipid channel (left) identifying LNP position and size and mRNA channel (middle) quantifying mRNA copy number, and merged image (right) shows colocalized lipids and cargo. Quantitative image analysis extracts nanoscale dimensions and cargo copy number per LNP. Scale bar, 10 µm. **d**, 2D histogram of the number of mRNA copies per LNP versus LNP size. Data are from single-particle LNP measurements as shown in **c**. *N,* 110,635. Step 2 (**a,e**–**f**). **e**, Internalized LNPs (magenta) are quantified using volumetric imaging in live cells stained with CellMask green (cyan), 1 h post-incubation and wash. Left: volumetric image. Right: 2D slice. Scale bars, 10 µm. **f**, Distribution of LNPs per cell. Mean ± SEM of three technical replicates. Step 3 (**a**,**g**–**i**)**. f**, EGFP expression in live transfected cells recorded using spinning disc confocal microscopy. Scale bar, 20 µm. **h**, Single cell from **g** at three timepoints shows time dependent increasing EGFP intensity. Segmentation masks (yellow) extracted fluorescence of each cell. Example mask shown (12 h). Scale bars, 5 µm. **i**, 3D plot of EGFP intensity of individual cells versus time. Green shading indicates mean intensity; orange line, mean trajectory. *N* shown, 9,563; *N* total, 95,637 across three technical replicates of data recorded every 30 min for downstream model evaluation. Model evaluation (**a**,**j**–**m**). Predicted release rates, the multiple acquisition timepoints in **i**, were used for determine prediction stability through 5,000 independent inference iterations based on 3 timepoints drawn from **i**, following the same “rules” as simulated data. **j,** Histogram of predictions from 3 technical replicates of the same assay (blue, yellow, green) and their average (orange) revealing a mean prediction of 7.82 ± 0.42%. Note, the model informed by experimental noise and cell-to-cell variability also outputs the full spectrum of outcomes rather than a single value. The histogram shows the results of the 5,000 independent inferences for each of the replicates along the mean across the replicates. (**k**–**m**) Lower-triangular heatmaps evaluate model sensitivity to measurement timing: horizontal axis denotes interval between first and second measurement; vertical axis denotes interval between second and third measurement. **k**, Mean predicted release rate. **l**, Relative standard deviation across replicates. **m**, Predicted uncertainty (standard deviation from log-variance). Predicted uncertainty (std from log-variance). Additional statistics were extracted, further supporting the shown heatmaps (Supplementary Fig. 8).

To validate performance, we used an LNP formulation containing SM-102, the widely adopted as a benchmark ionizable lipid in mRNA delivery (28,29) and cholesterol (LNP/chol). Implementation of 0.2 mol% ATTO655-labeled DOPE lipid ensures interoperability for LNP loading and internalization experiments (30,31). Using a net negative charge of the fluorophore prevents interaction with mRNA within the LNPs (32,33) to ensure that the single LNP fluorescence intensity scales with the LNP’s surface area. Transfection was performed on HEK293 cells due to their extensive use in research, their well-documented susceptibility to LNP transfection, and their widespread role in the development of LNP systems and mRNA therapeutics (34).

Quantification of the number of mRNAs packed into each nanoscale LNP was based on our single-particle assays (33,35,36). ATTO655-lipid and AlexaFluor-488–labeled EGFP-mRNA (AF488-mRNA) immobilized on passivated coverslips for TIRF microscopy (Fig. 3b,c). Current reports, which rely on average readouts, vary widely, with values ranging from ∼2 to ∼100 mRNAs per LNP for systems of comparable size (13,37–39). Each LNP appears as a diffraction limited spot, but quantitative analysis allows extraction of their nanoscale dimension (37–39). A key element is the implementation of the negatively charged membrane marker ATTO655-lipid that is not electrostatically attracted to mRNA (32). DLS studies confirm the size distribution (see Methods and Supplementary Fig. 9). Dual-channel Imaging and colocalization analysis revealed that 86.1 ± 2.0% of LNPs contained mRNA (Supplementary Fig. 10), consistent with literature values. (38,40). Quantification of the background-corrected AF488-mRNA signal, after calibration with individual AF488-mRNA (see Methods and Supplementary Fig. 11), provided the copy numbers of mRNA load per LNP and its dependence on LNP diameter. The single-particle readout revealed that LNPs contain an average of 6.82 ± 0.47 mRNA copies per LNP (Fig. 3d), consistent with current reports of ∼2 to ∼100 mRNAs per LNP for systems of comparable size (37–39).

To quantify LNP cellular uptake, live HEK293 cells stained with CellMask Green plasma membrane stain were incubated for 1 h with LNP/chol at a concentration of 500 ng/mL mRNA, washed, and imaged volumetrically using two-color lattice light-sheet microscopy (LLSM) (Fig. 3e). Note, alternative volumetric imaging microscopy techniques like SDCM or light-sheet etc. could be employed just as effectively for this step. Three-dimensional segmentation of the membrane channel generated per-cell masks from which internalized LNP fluorescence was extracted and converted into absolute particle counts using intensity distributions of individual LNPs (Fig. 3d and Supplementary Fig. 12) (1–4,13,41). Using this single-particle approach, we determined that HEK293 cells internalize on average 43.99 ± 2.17 LNPs per cell (Supplementary Fig. 13). These values are consistent with the expected uptake range and, to the best of our knowledge, provide the first direct numerical alignment of relative uptake with absolute intracellular particle numbers (42,43).

### Conversion of time dependent EGFP expression to mRNA release ratio

The third step for predicting the experimental release ratio involved acquiring temporal single-cell expression profiles of the fluorescent marker EGFP, along with the fourth meta-parameter, the total number of cells. Cells were seeded on conventional flat-bottomed slides to ensure system adaptability and minimize potential perturbation from micro-patterned slides. EGFP expression in live individual HEK293 cells transfected with LNP/chol was recorded using SDCM every 30 min for 48 h (Fig. 3g,h). This dense timepoint recording was chosen to evaluate the model’s stability and sensitivity to measurement timing, while prolonged imaging ensured that the entire process, including the decrease in EGFP signal due to protein degradation, was captured for model training.

Time-dependent parallelized recording and cell segmentation using Cellpose3 (44) (Fig. 3h, Supplementary Fig. 14–15) enabled extraction of fluorescence intensity for each of the more than 90,000 recorded cells across timepoints and replicates. We also quantified the fraction of cells that did not express any EGFP to address non-productive mRNA delivery (Supplementary Fig. 16). As expected, the recorded single-cell EGFP intensities increased during protein translation and subsequently decreased due to protein degradation (Supplementary Fig. 17), consistent with our simulation results (see Fig. 3i for 9,563 cells and Supplementary Fig. 18 for all data).

Quantification of release ratio by RNASCAPE averaged to 7.82 ± 0.42% for LNP/chol in HEK293 cells across three biological replicates (Fig. 3j). This value is notable as it is derived from non-fluorescently labeled mRNA on naïve cell line and yet aligns closely with independent experimental estimates reliant on specialized cell lines expressing specific markers (1–6), supporting both the model’s validity and the biological accuracy of the predicted release efficiency.

To evaluate the robustness of this prediction, we performed 5,000 independent inference iterations in which measurement timepoints were randomly resampled under the same temporal constraints as the simulations (see Methods). The distribution of predictions from individual iterations (Fig. 3j) is further explored in three lower-triangular heatmaps depicting the average predicted release ratio, relative standard deviation, and predicted uncertainty (Fig. 3k). Higher predicted release ratios were observed at the extremes of possible time separations between measurements (upper triangle region), corresponding to the histogram peak around 8.0%. Conversely, a trend toward lower predictions is evident in the bottom-left region of the heatmap, consistent with the small left-side distribution of the primary histogram peak.

Further analysis of inter-replicate variability (Fig. 3l) showed that measurement intervals of approximately 20 h between the first and second timepoints resulted in predictions with the least variability and highest consensus. This suggests inherent stability in the model’s predictions at these intermediate measurement intervals. The predicted uncertainty on the release ratio, expressed as the standard deviation derived from log-variance, mirrored these trends (Fig. 3k), with variance increasing modestly at higher release ratios. Thus, RNASCAPE provides stable and reproducible escape estimates when temporal sampling spans 15–25 h between measurements, allowing for easy adaptation across different experimental configurations and imaging schedules without requiring specialized hardware, precise timing, or specialized cell lines.

### RNASCAPE reveals sterol effect on LNP mRNA loading and functional delivery

To expand the operational utility of RNASCAPE, we utilized it to quantify the effect of LNP composition and cell line on mRNA release, and by utilizing different microscopy systems. We compared three different conditions: LNP/chol and SM-102/β-sitosterol LNPs (LNP/β-sito) in HEK293 cells and SM-102/β-sito in HeLa cells using volumetric LLSM as the imaging method.

The volumetric nature of LLSM introduces distinct illumination and detection geometries compared with SDCM, resulting in differences in signal-to-noise, spatial resolution, and temporal sampling (45,46). To further distinguish the datasets, each timepoint was acquired on a separate microscopy slide, ensuring that unique cells were imaged at every timepoint, contrary to the SDCM experiments, where the same cells were imaged continuously. Cells were imaged at 6 h, 21 h, and 44 h post-transfection, using 15 h and 23 h intervals selected from the optimal temporal spacing analysis (Fig. 3j–m). Because each LLSM timepoint was acquired on a separate slide, unique cells were measured at each timepoint. At each timepoint, cells were segmented in 3D from the EGFP channel, and single-cell expression intensity distributions were extracted for three technical replicates (Fig. 4a,b) and were input to RNASCAPE. To ensure convenient use of the model, we developed a fast, simple, and free-to-use graphical user interface (GUI) (Fig. 4c).

**Fig. 4:**
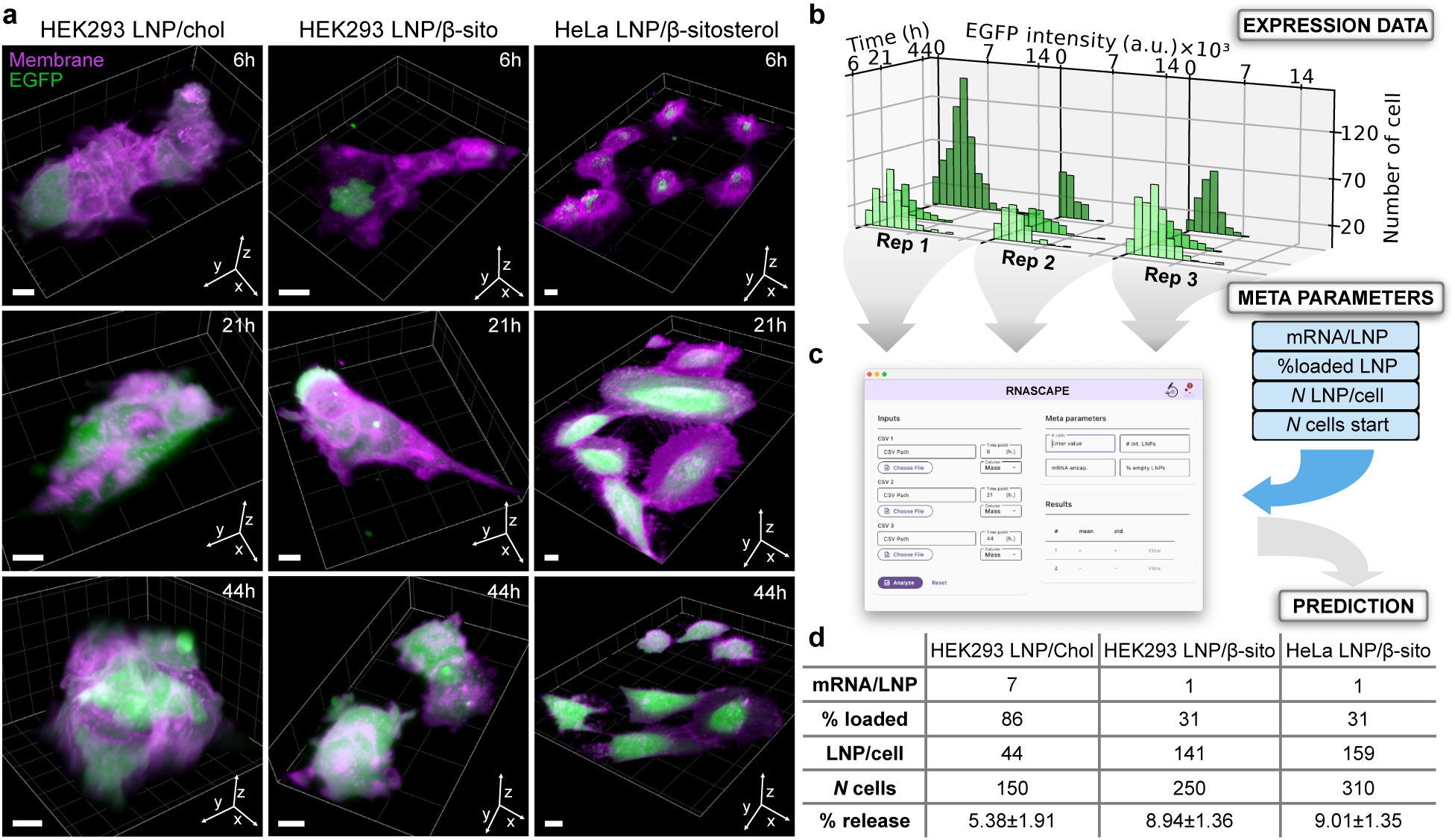
Comparison of mRNA release ratio of two LNP compositions and cell lines using LLSM expression data. **a**, Volumetric LLSM data of HEK293 and HeLa cells stained with CellMask deep red plasma membrane stain expressing EGFP transfected with transfected with LNP/chol and SM-102/β-sito at 6 h, 21 h, and 44 h after LNP. Note, image contrast is set differently for the two LNP formulations for visibility. Scale bars, 10 µm. **b**, Histograms of single cell EGFP expression distributions from segmented LNP/chol data presented in ***a***. Data display the expression profiles of three technical replicates at three timepoints. *N* rep 1, 927; *N* rep 2, 293; *N* rep 3, 632. **c**, Screenshot of the provided RNASCAPE standalone GUI that automates and simplifies analysis. The intuitive user interface allows users with only a few clicks to input the experimentally acquired replicates of EGFP expression data at three timepoints and the four experiment-specific meta-parameters. **d**, Meta parameters used as model input and predicted mRNA release ratio for two LNP compositions and two cell lines. Predictions show mean ± std. of three technical replicates.

Using RNASCAPE, we estimated mRNA release ratios from the three-timepoint expression distributions (Fig. 4d and Supplementary Fig. 19) using composition and cell line-specific meta-parameters (Supplementary Fig. 13 and Supplementary Fig. 20). Beyond differing by their sterol, the two LNP compositions differ by their mRNA packing where LNP/chol has 86% loaded LNPs with 7 mRNA per loaded LNP and LNP/β-sito has 31% loaded LNPs with 1 mRNA/loaded LNP. In HEK293, LNP/β-sito internalizes more than LNP/chol at 141 LNP per cell and 44 LNP per cell, respectively. However, between HEK293 and HeLa, LNP/β-sito internalizes more evenly with 159 LNPs per cell in HeLa. The number of analyzed cells was similar (150, 250, and 310 cells). Based on these inputs, RNASCAPE predicted release ratios of 5.38 ± 1.91% for LNP/col in HEK293 cells, which is similar to our validation prediction from SDCM, and predicted 8.94 ± 1.36% release for LNP/β-sito in HEK293 and 9.01 ± 1.35% in HeLa. The predicted release ratios are consistent with the SDCM validation range and published estimates, supporting the use of RNASCAPE for comparing LNP compositions across imaging setups.

## DISCUSSION

Endosomal escape is the critical bottleneck in mRNA delivery, yet the field lacks a standardized method to quantify it. Bulk reporter assays are simple but mask the readouts of uptake, escape, and translation. Imaging methods like smFISH provide mechanistic detail but are invasive, low throughput, and offer limited information on functional mRNA delivery. Galectin-9 assays, in addition to requiring specific stable cell lines, assume a one-to-one correlation between endosomal damage and escape. Analytical models, meanwhile, capture averages but miss single-cell variability. RNASCAPE addresses these limitations by delivering rapid, robust, and reproducible estimates of functional escape with minimal input, offering the first scalable framework for benchmarking LNP performance.

A defining strength of RNASCAPE is its robustness and generalizability. We validated performance across spinning disc confocal microscopy and lattice light-sheet microscopy, two platforms with very different optical geometries, sampling strategies, and noise properties. The framework does not depend on the modality itself, but on the structure of the input data consisting of three timepoints of distributional expression combined with four easily measurable meta-parameters. This structure makes RNASCAPE inherently adaptable across systems. With retraining, the same pipeline could be applied to alternative readouts such as flow cytometry-based measurements. Likewise, it can be extended across cell types and LNP formulations. In this way, RNASCAPE provides not just a model but a framework for functional escape quantification.

In our comparison of LNP compositions and cell lines, RNASCAPE predicted a two-fold difference in release between LNP/chol and LNP/β-sito in HEK293 cells but near identical release for LNP/β-sito in HEK293 and HeLa cells, revealing clear formulation-specific release that is independent of the chosen cell lines.

The absence of reliable standardized quantification methods of mRNA escape has left endosomal escape studies fragmented, with results often difficult to compare across laboratories. Community-driven tools in other areas such as CellProfiler in imaging (47), SCANPY in single-cell transcriptomics (48), or benchmark platforms in drug screening (49) have shown how open and reproducible frameworks can accelerate discovery and create common reference points. RNASCAPE has the potential to provide the same function for nucleic acid delivery, offering convenient and reproducible readouts of functional escape. RNASCAPE implemented in screening readouts can provide a complex fingerprint of correlations of lipids to RNA and cell type escape benchmarking the field across laboratories industrial stakeholders and formulations. This can transform isolated studies into libraries of comparable fingerprints that can be utilized to inform the rational design of rational design of formulations, either nucleic acid modifications, lipid composition, or N/P ratio with improved functional mRNA escape.

To facilitate this and ease experimental use and adoption in laboratories without computational expertise, we designed RNASCAPE as a standalone GUI. With only a few clicks, users can upload their EGFP expression distributions and four meta-parameters: average LNPs per cell, average mRNA per LNP, fraction of loaded particles, and number of cells at the start of the EGFP expression measurements. The software then automatically outputs predicted release ratios and generates distributional outputs. Its modular design allows users to run the entire workflow from start to finish or focus on individual steps such as meta-parameter calibration or fingerprint extraction. The open-source implementation, on the other hand, allows expert users to extend or adapt the pipeline, for example to include new reporter systems or to explore parameter sensitivity. The combination of simplicity for broad use and flexibility for customization allows convenient implementation of RNASCAPE across the community for pipelining, screening, and benchmarking quantifying endosomal escape.

Our approach to quantifying functional mRNA release relies on the time-dependent increase in downstream fluorescent reporter protein densities in cells, supported by simulations spanning biologically and experimentally realistic parameter ranges. Notably, no experimental method currently provides a direct measurement of the absolute cytosolic escape fraction of delivered cargo, reflecting the inherent complexity of intracellular trafficking and release. As a result, the validity of RNASCAPE rests on the biophysical assumptions encoded in the simulation framework. While unmodeled sources of biological heterogeneity may limit direct correspondence to any single experimental ground truth, the model establishes a principled and internally consistent baseline, enabling quantitative comparison across conditions, formulations, and cell types. In this context, accuracy should be interpreted as functional consistency across experiments. To capture experimentally relevant heterogeneity, we used 10–1000 cells per experiment, balancing biological plausibility with computational compliance. While the current dataset includes 800,000 simulated experiments, each defined by thirteen kinetic parameters, it can be further extended given sufficient computational resources. The method outputs mRNA escape for the EGFP fluorescent reporter, and its adaptability allows evaluation of whether mRNAs encoding additional fluorescent reporters exhibit similar escape fractions.

Our predictions converge on escape efficiencies of 5–9%, consistent with the dominant estimates in the literature and reproducible across replicates and imaging modalities. The occasional bimodality in distributions should not be regarded as noise but rather as a reflection of the stochastic nature of escape and the sensitivity of predictions to timing. Such multimodal outcomes may contain biologically meaningful signatures of distinct escape pathways or cellular states resulting in distinct escape rates. Limitations remain, however, as our predictions are indirect, relying on EGFP expression rather than direct RNA counts, and the transformer-based architecture cannot yield mechanistic rate constants, that is, it predicts functional outcomes rather than explicit kinetic parameters describing processes like uptake speed, endosomal rupture frequency, or mRNA decay. These limitations highlight the complementarity with existing quantitative methods as imaging and kinetic models provide mechanistic clarity, while RNASCAPE provides the functional number. This approach may enable the field to compare formulations, conditions, and strategies based on a functional metric. By embedding mechanistic realism in simulations, predictive power in machine learning, and accessibility in a GUI, RNASCAPE takes a step toward community-driven reproducibility and progress in overcoming one of the most persistent challenges in nucleic acid delivery.

## MATERIALS AND METHODS

### Materials

ATTO655-DOPE was purchased from ATTO-TEC (Siegen, Germany). DSPC, DMG-PEG2000, and SM-102 lipids were purchased from Avanti Polar Lipids Inc. CleanCap® EGFP mRNA (5moU) was purchased from TriLink Biotechnologies. DMEM - high glucose, FBS, Triton X100, Quant-it™ RiboGreen assay, CellMask green plasma membrane stain, CellMask deep red plasma membrane stain, L-Glu, pyruvate, DMSO, cycloheximide, DPBS were purchased from Sigma-Aldrich Denmark. Citrate, DEPC treated water, PBS from Invitrogen^TM^. DMEM without phenol red, Ethanol 99%+, NeuTravidin, TetraSpeck™ Microspheres, 0.1 μm from Fisher Scientific^TM^. AF488-EGFP-mRNA was purchaced from RIBOPRO. PLL-g-PEG and PLL-g-PEG-biotin were purchased from SuSoS AG. All microscopy slides and accesories were purchased from ibidi. Cells were obtained from ATCC.

### Experimental procedures LNP preparation

LNPs were prepared using a custom microfluidics system fitted with a glass herringbone mixer (Darwin Microfluidics) by mixing at a flow rate ratio of 3:1 (aqueous:organic) with the aqueous phase consisting of mRNA in dissolved in in RNase-free 10 mM citrate buffer pH 4, and the organic phase consisting of lipids dissolved in 99%+ ethanol. LNPs containing SM-102 and cholesterol were formulated at a total flow rate of 500 µL/min. Lipids were prepared in at molar ratios (50:10:38.3:1.5:0.2 for SM-102: DSPC: cholesterol: DMG-PEG2000: ATTO655-DOPE) at a total lipid concentration of 1.2 mg/mL. LNPs containing SM-102 and β-sitosterol were formulated at a total flow rate of 100 µL/min. Lipids were prepared at molar ratios (50:9.5:38.5:1.0:0.5 for SM-102: DSPC: β-sitosterol: DMG-PEG2000: ATTO655-DOPE) at a total lipid concentration of 5.0 mg/mL. The mRNA cargo consisted of CleanCap® EGFP mRNA with full 5moU substitutions. mRNA was used in formulation at a N/P ratio of 6. Immediately following formulation, ethanol was removed through diafiltration by two consecutive rounds of diluting samples in cold PBS and centrifugation at 3000 × *g* in 4mL 10 kDa MWCO Amicon® Ultra centrifugal filters (Millipore) at 4°C before storage at 4°C.

SM-102/cholesterol LNPs for single-particle TIRF microscopy were prepared as above with molar ratios (50:10:38.3:1.0:0.5:0.2 for SM-102: DSPC: cholesterol: DMG-PEG2000: DSPE-PEG2000-biotin: ATTO655-DOPE) and AF488-EGFP-mRNA. SM-102/β-sitosterol LNPs for single-particle TIRF microscopy were prepared as above with molar ratios (50:9.5:38.5:0.5:1.0:0.5 for SM-102: DSPC: β-sitosterol: DSPE-PEG2000-biotin: DMG-PEG2000: ATTO655-DOPE) and AF488-EGFP-mRNA.

### LNP characterization

The size and polydispersity of LNPs was measured using a Malvern Panalytical Zetaziser DLS machine. LNPs were diluted in PBS and measured in triplicates at 25°C using a built-in fluorescence filter to accommodate ATTO655-DOPE in the LNPs.

Total mRNA concentration was measured on a NanoDrop 2000 spectrophotometer from Thermo Scientific. Encapsulation efficiency (EE%) of mRNA in LNPs was measured using the Quant-it™ RiboGreen assay. Here, the amount of unencapsulated mRNA amount is measured on intact LNPs while the total mRNA amount is measured on LNP disrupted by pre-incubation with 0.1% v/v Triton X-100. EE% was calculated as:

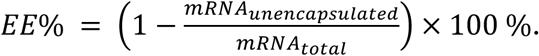

### Single-particle LNP characterization using TIRF microscopy

Single-particle characterization of LNP mRNA loading was performed as described previously (33) with modifications use a 640 nm excitation wavelength laser for imaging ATTO655-labeled lipids and a 488 nm excitation wavelength laser to image ATTO488-labeled mRNA.

### Cell culturing

HEK293 (ATCC, cat: CRL-1573) HeLa (ATCC, cat: CCL-2) cells were maintained in DMEM supplemented with 10% FBS at 37°C, 5% CO2, and 70% humidity and were passaged every 3 to 5 days. All experiments on HEK293 cells were performed on passage number 7 through 10, while all experiments on HeLa cells were performed on passage number 11 through 13.

### Preparation of cells for microscopy

In preparation for microscopy experiments, cells were seeded at 5,000 cells per well in ibidi ibiTreat 8-well µ-slides for SDCM experiments, or ibidi glass-bottom 8-well µ-slides for TIRF and LLSM experiments, and incubated at 37°C, 5% CO2, and 70% humidity for 1 day.

For SDCM experiments, cells were washed once in DPBS before addition of imaging medium containing LNPs at a concentration of 500 ng/mL mRNA and incubating for 1 h. Imaging medium containing LNP was then aspirateda nd cells washed once in DPBS before adding imaging medium without LNP.

For LLSM experiments of EGFP expression at 3 timepoints, all cells were seeded at the same time and were at either 6, 21 or 44 h prior to imaging washed once in DPBS and incubated with imaging medium containing LNPs 500 ng/mL mRNA for 1 h before washing once in DPBS and incubating finally incubating in imaging medium. Prior to imaging, cells were stained for 5 minutes with 1:1000 dilution of CellMask deep red in imaging medium before the final transfer to imaging medium.

For experiments investigating the internalization of LNPs in cells on LLSM, cells were washed once in DPBS before being incubated with imaging medium containing LNPs 500 ng/mL mRNA for 1 h or a control medium without LNP for 1 h. Cells were then washed and stained for 5 minutes with 1:000 dilution of CellMask green plasma membrane stain in imaging medium before being washed once in DPBS and imaged in imaging medium.

To estimate the degradation rate of EGFP in live cells on SDCM, cells were incubated with imaging medium containing LNPs 500 ng/mL mRNA for 1 h before washing in DPBS and incubation in imaging medium for at 37°C, 5% CO2, and 70% humidity for 24 h. The medium was then aspirated and replaced with imaging medium containing 50 µg/mL cycloheximide before starting image acquisition.

### Live-cell imaging across platforms

#### Spinning disc confocal microscopy

All experiments of continuous expression or degradation of EGFP in live HEK293 cells were conducted on an inverted spinning disc confocal microscope (SDCM) (Olympus SpinSR10) using a 20X magnification air objective (Olympus) with an NA of 0.50 and a CMOS camera (photometrics PRIME 95B) at 37°C, 5% CO2, and 70% humidity. The contour of cells was imaged using LED brightfield illumination lamp at 50.04 ms exposure time while expressed EGFP was excited using a 488 nm laser line at 20% laser power and imaged with 50.04 ms exposure time. For each experiment, 5 to 30 positions were selected in the microscopy slide well based on the distribution and singularity of cells. Each position was imaged by recording a *z*-stack consisting of 7 slices spaced 3 µm with the center slide position near the microscopy slide surface. Starting 30 minutes after incubation with LNP, all positions were imaged every 30 minutes for a total of 48 h. EGFP expression experiments were performed in 3 technical replicates. The resulting pixel size was 0.917 × 0.917 µm.

#### Lattice light-sheet microscopy

All LLSM experiments were performed on a ZEISS Lattice Light-sheet 7 equipped with Hamamatsu ORCA-FusionBT CMOS cameras. Data was acquired using a 30 × 1000 µm light-sheet cropped to a width of 296.94 µm and height of 55.532 µm for experiments involving HEK293 cells and 21.604 µm for experiments involving HeLa cells, and an inter-frame step size of 200 nm. Prior to data acquisition, the light-sheet was aligned by measuring the volumetric point spread function (PSF) of 0.1 µm TetraSpeck™ Microspheres diluted 1:100 in high gel agarose and cast in an empty well on the same microscopy slide as the cell sample. Voxel size 0.145 × 0.145 × 0.145 µm

#### Volumetric measurements of EGFP expression on LLSM

Cells were incubated with LNP for 1 h and then left to express EGFP for 6, 21, or 44 h before microscopy experiments. EGFP was excited using a 488 nm laser channel at 80% laser power and 5.0 ms exposure time while CellMask Deep Red was excited using a 640 nm laser channelat 5% laser power and 5.0 ms exposure time. Each timepoint was recorded in three technical replicates containing 5 to 7 fields of view per replicate. HEK293 LNP/chol experiments were recorded using a 500–548 nm and a 575–618 nm bandpass filter set on the emission of the 488 nm laser channel and a 659–720 nm bandpass filter on the 640 nm laser channel. HEK293 LNP/β-sito and HeLa LNP/β-sito were both recorded using a 500–548 nm and a 659–720 nm bandpass filter set on the emission of the 488 nm laser channel.

#### Single LNP characterization on LLSM

LNPs were diluted in PBS to 50 ng/mL mRNA and added to ibidi glass-bottom 8-well µ slides for 30 minutes to settle spontaneously on the slide surface. The glass surface was then positioned in the middle of the field of view, and LNPs were imaged using a 640 nm laser line at 40% laser power and 10.0 ms exposure time. 1000 slices were recorded in three technical replicates. LNP/chol experiments were recorded using a 500–548 nm and a 575–618 nm bandpass filter set on the emission of the 488 nm laser channel. LNP/β-sito were recorded using a 500–548 nm and a 659–720 nm bandpass filter set on the emission of the 488 nm laser channel.

#### LNP internalization measurements on LLSM

After incubation with LNP or control medium and staining with CellMask green plasma membrane stain, cells were imaged by exciting the LNPs with a 640 nm laser channel at 40% laser power and 10.0 ms exposure time while the membrane stain was imaged using a 488 nm laser channel at 4.0% laser power and 10.0 ms exposure time. Each condition was recorded in three technical replicates containing 7 to 8 fields of view per replicate. HEK293 LNP/chol experiments were recorded using a 500–548 nm and a 575–618 nm bandpass filter set on the emission of the 488 nm laser channel and a 659–720 nm bandpass filter on the 640 nm laser channel. HEK293 LNP/β-sito and HeLa LNP/β-sito were both recorded using a 500–548 nm and a 659–720 nm bandpass filter set on the emission of the 488 nm laser channel.

### Analytical methods

#### TIRF microscopy analysis

Quantification of LNP size and mRNA loading was performed as described previously (33).

#### Spinning disc confocal microscopy analysis

##### Selection of surface slice

Time-resolved EGFP expression data from SDCM was preprocessed to only perform analysis on *z*-planes recorded nearest to the microscopy slide surface. This was done by selecting the *z*-plane with the highest mean intensity at every timepoint in every position using python code.

##### Brightfield contrast enhancement

To enhance the contrast of each frame in a 7-slice image stack, we applied a two-step correction process. First, each 2D slice was normalized and rescaled to map intensity values to the range [0, 255], with saturation at the 98th percentile to suppress high-intensity outliers and enhance overall dynamic range using the scikit-image package. This first step was used as a manual clipping for Contrast Limited Adaptive Histogram Equalization (CLAHE) which was applied using OpenCV, allowing for locally adaptive contrast enhancement while preventing over-amplification of noise (50). The final corrected image was obtained by averaging the enhanced slices, resulting in a single, contrast-enhanced image representative of the stack.

#### Brightfield illumination profile correction

To account for the inherent radial illumination profile in the recorded brightfield data, an illumination profile correction python script was used. Any given raw image was blurred by a big Gaussian convolution (σ>=30 pixels) making the profile more dominant than the existing objects and fitted by a 2D Gaussian function. The 2D fitted Gaussian was normalized to its highest value and the raw image was divided by the normalized fitted Gaussian.

#### 2D cell segmentation

SDCM data were segmented using Cellpose3. An in-house model was created using a training set containing 50 fields of view of EGFP expression only in cells at various timepoints during expression to cover varying levels of EGFP intensity. Here, each image was first segmented using the built-in Cyto3 model with an initial segmentation diameter of 90 pixels model and the results were manually inspected and corrected in the Cellpose3 GUI before training the final model on the corrected masks. All SDCM data of EGFP expression were then segmented in the EGFP channel using the trained model with a segmentation diameter of 70 pixels and Cellpose3 in Python. Segmentation artefacts were filtered out from the final data by excluding masks having the 1% largest and 1% smallest area (exclusion of multiple cells in the same masks, senescent or multinucleated cells, cell debris etc.), the 1% highest variance in EGFP intensity (exclusion of multiple cells in the same mask or inclusion of background signal in masks), or eccentricity ≥ to 0.1 (exclusion of dead or mitotic cells).

### EGFP intensity extraction

The EGFP intensity in individual cells was then extracted by calculating the numerical average of the EGFP channel within individual masked cell areas given by Cellpose segmentation at all timepoints.

### Lattice light-sheet microscopy analysis

#### 3D particle detection and quantification of PSF

To perform 3D detections of the point spread function (PSF), the volumetric dataset was treated as a time-lapse sequence, with each z-slice corresponding to a single frame. Two-dimensional particle detection was carried out independently on each z-slice using a Laplacian of Gaussian (LoG) filter. The detected 2D particles were subsequently linked across adjacent z-slices using a nearest-neighbor algorithm to reconstruct 3D particle trajectories. Reconstructed 3D particles spanning fewer than five consecutive z-slices were excluded from further analysis to reduce the impact of spurious detections. For each remaining particle, the background-corrected intensity was calculated on a per-frame basis by subtracting the local background intensity from the detected signal. These intensity profiles were then fitted with a one-dimensional Gaussian function using chi-square minimization and the Minuit optimization (James1975). The Gaussian fitting procedure provided both the z-center of each 3D particle and a quantitative criterion for validating true positive detections. The total background-corrected intensity of each 3D particle was computed as the sum of the background-corrected intensities across all linked frames.

Intensity values from all fields of view within each replicate were pooled, and the characteristic particle intensity was estimated as the mode of the resulting distribution using Gaussian kernel density estimation in linear intensity space. For datasets with three replicates, replicate-specific modes were averaged and the standard deviation across replicates was reported.

#### 3D cell segmentation

Volumetric segmentation of cells imaged with LLSM was performed using a hybrid approach combining Cellpose3 (44) and the Segment Anything Model 2 (SAM 2) (52). We adapted SAM 2 video segmentation capabilities to propagate 2D segmentations across z-stacks, treating sequential z-slices analogously to video frames. For each recorded volume, we first identified an optimal reference slice, typically near the cell’s mid-height where membrane contrast and cell boundaries were most clearly defined. This reference slice from the membrane channel was segmented using Cellpose3 (Cyto3 model) and manually refined in the Cellpose GUI to ensure accurate seed masks. To extend segmentation across the full z-stack, we split the volume at the reference slice into two sub-stacks: one extending toward lower z-values (z = 0) and one toward higher z-values (z = max). Using the centroid of each cell mask as an initial prompt, SAM 2 propagated the segmentation bidirectionally through each sub-stack, tracking cells across sequential slices. The two segmented sub-stacks were then merged to produce complete 3D cell volumes. This approach leverages the robust 2D segmentation of Cellpose for initialization while exploiting SAM 2 temporal consistency mechanisms for efficient and accurate 3D propagation.

### EGFP intensity extraction

From the 3D cell segmentations, the EGFP intensity values of all voxels contained within individual mask were summed and normalized to the cell volume to provide a mean EGFP intensity per cell.

### LNP internalized intensity extraction

To accurately measure internalized LNP signal (640 nm channel), we first corrected for background and cellular autofluorescence using untransfected control cells. The mean voxel intensity across all segmented untransfected cell volumes established a baseline background level. For transfected cells, we calculated the mean voxel intensity within each cell mask and subtracted the background value to isolate the true LNP-specific signal. This background-corrected intensity was then multiplied by cell volume to obtain the total integrated LNP signal per cell. Finally, by dividing this total cellular LNP intensity by the mean intensity of a single LNP particle (measured independently), we estimated the absolute number of internalized LNPs per cell.

### Particle quantification

To quantify internalized particles, 3D segmented per-cell masks were applied to the LNP channel to extract intracellular fluorescence, excluding extracellular and surface-bound particles. Calibration was performed using surface-tethered LNPs imaged under identical acquisition conditions, where volumetric integration of fluorescence defined the intensity of a single particle. Intracellular LNP signal per cell was then converted to absolute particle counts by dividing by this single-particle intensity. Integrating signal across the full cell volume, rather than detecting individual puncta, ensured that LNP–endosome fusion events within the 1 h incubation were captured, as degraded or fused particles may no longer appear spot-like.

### Analytical model of protein expression in cells

#### Parameters

- 𝑡 : evaluation timepoint
- 𝐼_0_ : initial number of internalized LNPs
- 𝑒_𝑟_ : fraction of LNPs that do not contain mRNA
- 𝑟 : probability of mRNA release from LNP
- 𝜆 : average mRNA loading per non-empty LNP, assuming Poisson distribution
- G: scaling factor representing the amount of protein produced per mRNA per time unit
- 𝜏_𝑚𝑅𝑁𝐴_ : half-life of mRNA
- 𝛾_𝑚𝑅𝑁𝐴_ : degradation rate of mRNA, where 𝛾_𝑚𝑅𝑁𝐴_ = 𝑙𝑛(2)/𝜏_𝑚𝑅𝑁𝐴_
- 𝜏_𝑝𝑟𝑜𝑡𝑒𝑖𝑛_ : half-life of protein
- 𝛾_𝑝𝑟𝑜𝑡𝑒𝑖𝑛_ : degradation rate of protein, where 𝛾_𝑝𝑟𝑜𝑡𝑒𝑖𝑛_ = 𝑙𝑛(2)/𝜏_𝑝𝑟𝑜𝑡𝑒𝑖𝑛_
- 𝑇_𝐷_: doubling time of cells
- 𝑣 : cell division rate, where 𝑣 = 𝑙𝑛(2)/𝑇_𝐷_
- 𝑁_𝑡_ : number of cells at time 𝑡

*<u>Cell division:</u>* The growth of cells is modeled assuming constant exponential division, assuming cells divide at a constant rate. The number of cells at time is given by equation 1.

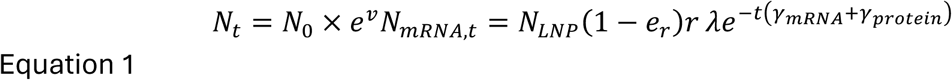

*<u>Average number of mRNA over time:</u> To estimate the concentration of mRNA at time t we consider that a fraction of LNPs do not contain mRNA, non-empty LNPs contain a, by a Poisson distributed, number of mRNAs, the encapsulated mRNAs release to the cytosol with certain probability, and the mRNA degrades exponentially. Additionally, cell division contributes to the dilution of the average mRNA count. The average mRNA count per cell is given by:*

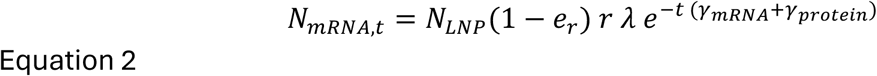

*<u>Average number of proteins over time:</u>* The estimated count at time t, is modeled based on mRNA availability, protein production, and the degradation rates of both mRNA and EGFP. These dynamics are governed by a constant term (the initial contribution), and a dynamic term accounting for degradation, which combine in equation 3.

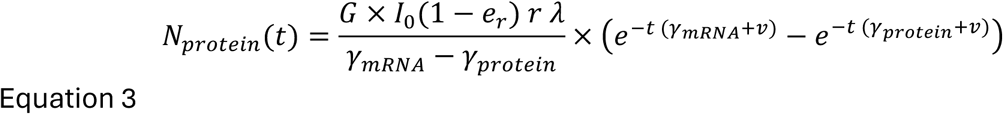

### Simulation of protein expression in cells

For a defined number of cells, the number of mRNAs released to the cytosol is first simulated. Each cell is assigned a diameter drawn from a lognormal distribution centered around the average cell size (σ = 0.2). The expected number of internalized LNPs per cell scales with the cell area, and a multiplicative lognormal noise term is applied (σ = 0.1) to introduce biological variability in uptake efficiency. The actual number of internalized LNPs per cell is then sampled from a Poisson distribution with this noisy, size-dependent mean. Among these LNPs, a specified fraction are empty, while the remaining non-empty LNPs each contain a number of mRNA molecules drawn from a Poisson distribution with a given average value (λ). Each contained mRNA has a fixed probability of being released to the cytosol. Together, this produces a distribution of released mRNA counts per cell that captures both biological noise and size-dependent uptake variation.

For each released mRNA, a degradation time is then simulated according to its half-life, corresponding to a constant probability of decay per time step. Each surviving mRNA expresses proteins at a constant rate until degradation, while every produced protein also has a probability of decay determined by its half-life. The time-resolved protein level is therefore determined by the cumulative balance between new protein production and degradation over time.

Cell division is simulated independently using a geometric distribution that defines the timing of each cell’s division events, based on the expected doubling time. At each division, all intact mRNAs and proteins are partitioned between the mother and daughter cell in a stochastic 50:50 manner. Transferred molecules retain their original degradation times, while new daughter cells begin with their inherited molecules and continue expressing and degrading proteins according to the same rules as their parents. Daughter cells are assigned new lognormally distributed sizes, but no new uptake of LNPs occurs after the initial time point. The simulation proceeds over successive generations until the final simulation time is reached.

This framework allows the estimation of the number of proteins in each cell over time, providing a means to compare simulated distributions with experimental data.

### Distributional fingerprint extraction

The distributional fingerprint constitutes 16 features for each time point, extracted from its corresponding EGFP expression distribution. The features aim to capture the general underlying trends in the distributions, allowing the ML model to elucidate the incremental changes over time.

### Machine learning models Architecture

The model takes 16 x 3 + 4 features as input, 16 features for 3 timepoints and 4 meta-parameters (Initial cell count, average internalized LNPs, ratio of empty LNPs, and LNP cargo loading efficiency). 15×3 features (excluding the “time from experiment starts”) as well as the meta-parameters are enriched through a 2-layer MLP, while the ‘time from experiment start’ feature is used to map a temporal encoding through a Time2Vec architecture. The temporal encoding features are linearly projected into the enriched features, which in turn are concatenated with the enriched meta-parameters along the sequence axis. A CLS token is appended to the resulting latent representation, and a learnable positional encoding is added before being further enriched through a multi-layered Transformer Encoder. The resulting position of the CLS token is then extracted and used through two MLP prediction heads that predict the release ratio and log-variance. Throughout the model, ReLU activation functions are used in the layer junctions, and a pre-layer-norm is applied in the transformer layers.

### Training

The model was trained for 400 epochs on an Apple M3 Max processor (MacBook Pro, 2023) using the AdamW optimizer (initial learning rate = 1 × 10⁻³). A *ReduceLROnPlateau* scheduler (patience = 10, factor = 0.8) was applied to adapt the learning rate during training. The loss function was a heteroscedastic regression loss, aligned with the model’s multi-output architecture. The simulated dataset (∼840,000 entries) was randomly partitioned into training (80%), validation (10%), and test (10%) subsets. The standardization values were calculated for the training set and used for the test and validation dataset for both the features, meta-parameters, and the release ratio (target). For further regularization, dropout was applied throughout the model, 20% for the initial enrichments MLPs as well as the linear projection, 50% for the transformer encoder and 40% for the prediction heads.

### Dataset

The training dataset was generated through simulations (see methods) in which parameters were uniformly sampled across a broad range, covering the diverse combinatorial systems listed in Table 1. For each drawn parameter set, three independent simulations were performed, one for each of the three extraction timepoints, to ensure independence of the resulting expression distributions. This approach avoided bias arising from the simulator’s implicit tracking of the same cells over time, which cannot be guaranteed in experimental measurements. Given the factorial combination of the different parameters, a big dataset was required, ∼840000 experimental triplets were simulated covering a stochastically spread grid across the parametric space. Further evaluation of this sufficiency is proven through the respectable performance on the test set, as parametric-set duplication is highly unlikely. For the preprocessing/feature extraction, 3 random timepoints were selected for each given simulation. The timepoints were selected in a manner such that the first timepoint always is before the 7^th^ h, and every consecutive timepoints will have at least 6 h of spacing between them. This method allows the model to generalize different timepoints and delta times between measurements. The features were extracted in a pair-wise manner, in such way that first timepoint is used for the first triplet, second timepoint for second triplet and third for third. The featurescollected as a single data entry along with its simulation parameters in a collective pre-processed dataset for training and validation.

## Authorship contributions

*Conceptualization*, F.H.S. and N.S.H. *Experimental design*, F.H.S. and N.S.H. *Experimental work*, F.H.S.: LNP formulation characterization of size and mRNA loading, time-resolved SDCM; F.H.S, G.B. and S.A.: LLSM of EGFP expression, LLSM of LNP internalization; A.B. and G.K.: LNP mRNA packing experiments; *Methodology*, F.H.S., T.O. and K.T.: Cell segmentation and LNP internalization analysis; K.T. and T.O.: LNP detection; F.H.S. and K.T.: EGFP intensity extraction; E.W.S.: Simulation conceptualization; E.W.S. with inputs of S.W.B.: simulation execution; E.W.S. and S.W.B.: statistical analysis and model convergence to data: E.W.S.: ML model conceptualization development of proof of concept models and curation of training data, S.W.B., with inputs of E.W.S.: Implementation of final ML model; M.W.D.: LNP mRNA packing analysis; F.H.S., E.W.S., A.B. and N.S.H.: control experiment design based on analysis. *Visualization*, F.H.S., E.W.S., and S.W.B. *Writing – original draft,* F.H.S., E.W.S., S.W.B., and N.S.H. *Writing – review and editing*, F.H.S., E.W.S., S.W.B., N.S.H., A.B., T.O., K.T., G.N. and M.W.D. *Supervision*: N.S.H. *Funding acquisition* N.S.H. and G.N., *Overall project management*, N.S.H.

## Acknowledgements

The work was financially supported by the Novo Nordisk Foundation via the Grand Challenge Programs in support of the Center for Optimized Oligo Escape and Control of Disease (NNF23OC0081287), and the Center for 4D cellular dynamics (NNF22OC0075851). The work was also financially supported by the Swiss National Science

Foundation (310030M_204518) and Villum Foundation Synergy grant (40578). The funding bodies were not involved in the work.

