## Supplementary Information for "Rapid and reliable quantification of cytosolic mRNA escape (RNASCAPE)"

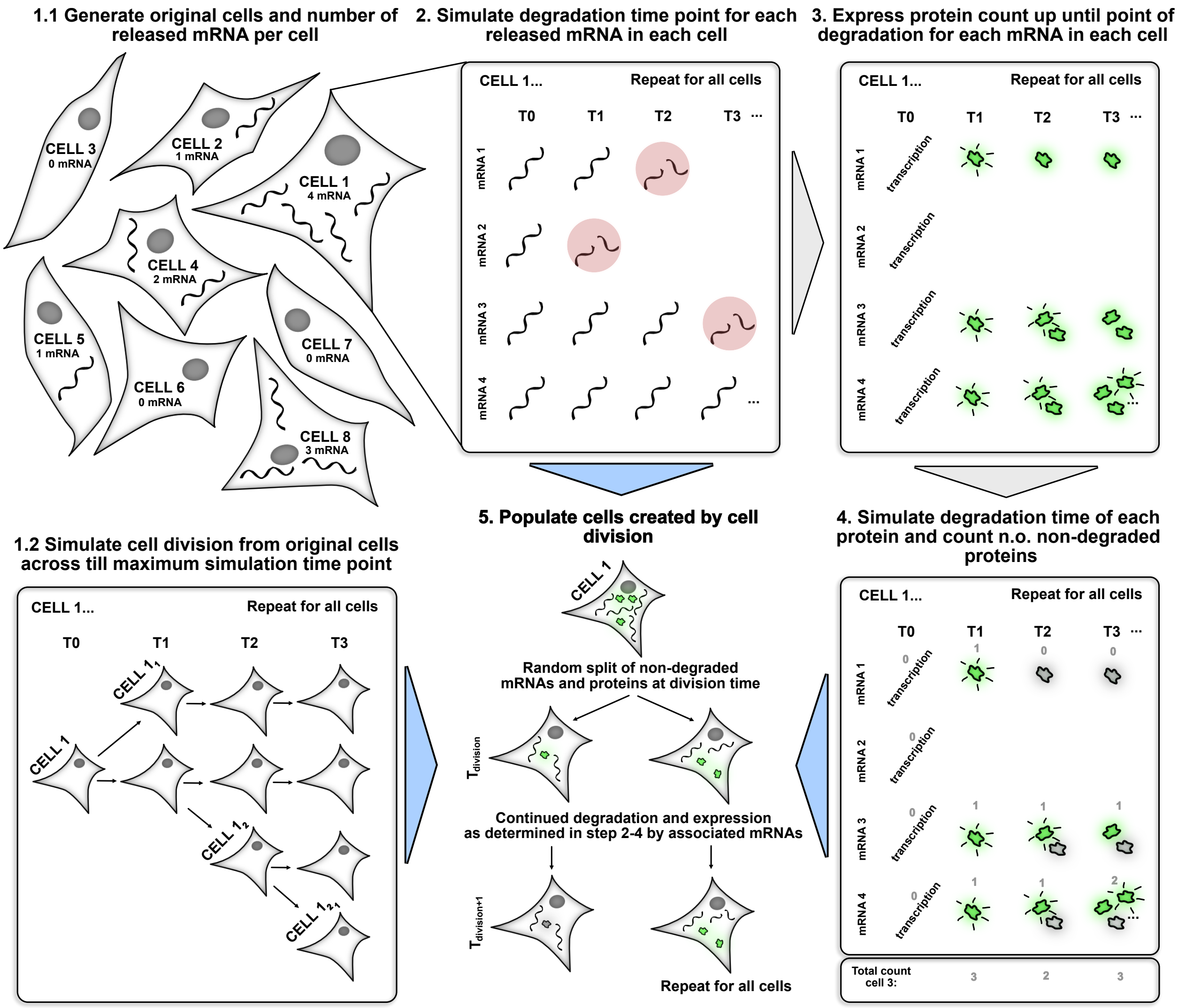


**Supplementary Fig. 1: Schematic simulation workflow.** The original / start number of cells is simulated as well as the number of mRNA released to each cell (1.1). The cell division is simultaneously simulated from time zero to the end time point of the simulation (1.2). For each mRNA released in 1.1 its degradation timepoint is determined (2). Hereafter, the number of proteins expressed for each mRNA (for each timepoint before its degradation) is determined (3). Lastly, the degradation timepoint for each of the expressed proteins is determined/simulated (4). Combining all the aforementioned steps, the mRNA and protein are split across the cells upon cell division. All expressed and non-degraded mRNA and protein is split evenly at the given timepoint of the cell division (5).

OAT: One at a time

PPS: Point-wise Perturbation Sensitivity (Error from baseline predictions)

APE: Absolute Percentage Error (Error from Ground Truth)

MAPE: Mean APE

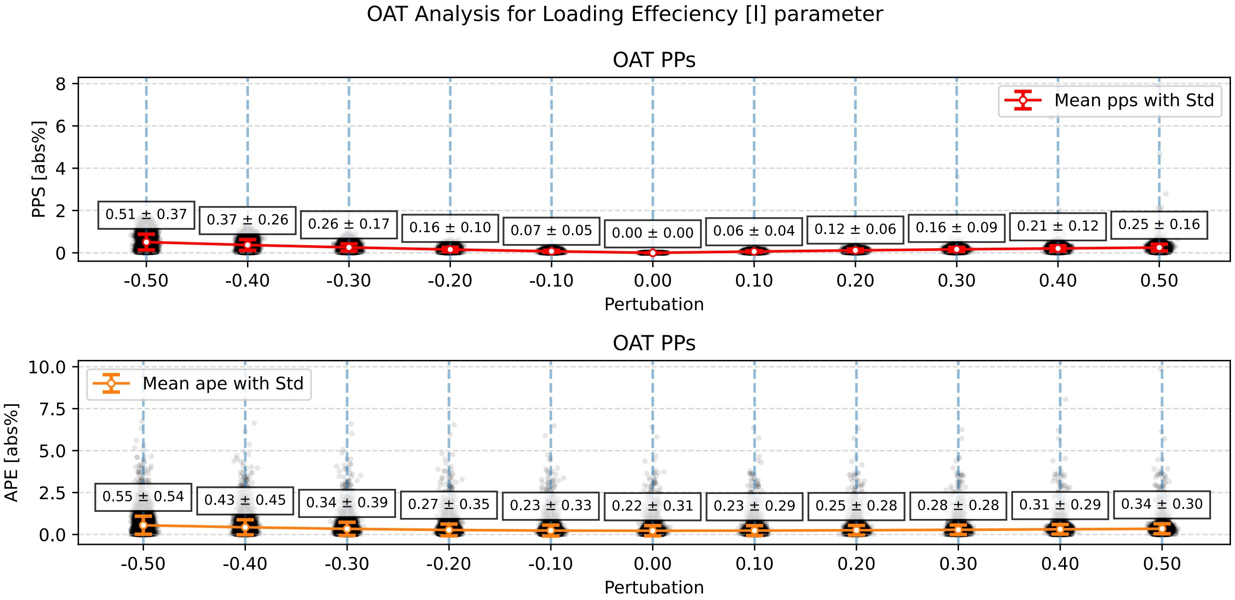


**Supplementary Fig 2: Prediction dependence on mRNA encapsulation through parameter perturbation. Top row: shows** One at a time (OAT) for the point wise perturbation sensitivity (PPS), a trend can be seen as the perturbation increases, with the strongest effect at the 50% negative perturbation at a relative change of 50%. Bottom row: Shows the Absolute Percentage Error (APE) from the simulated test set. The same behavior can be seen here however, reflected in the average relative error.

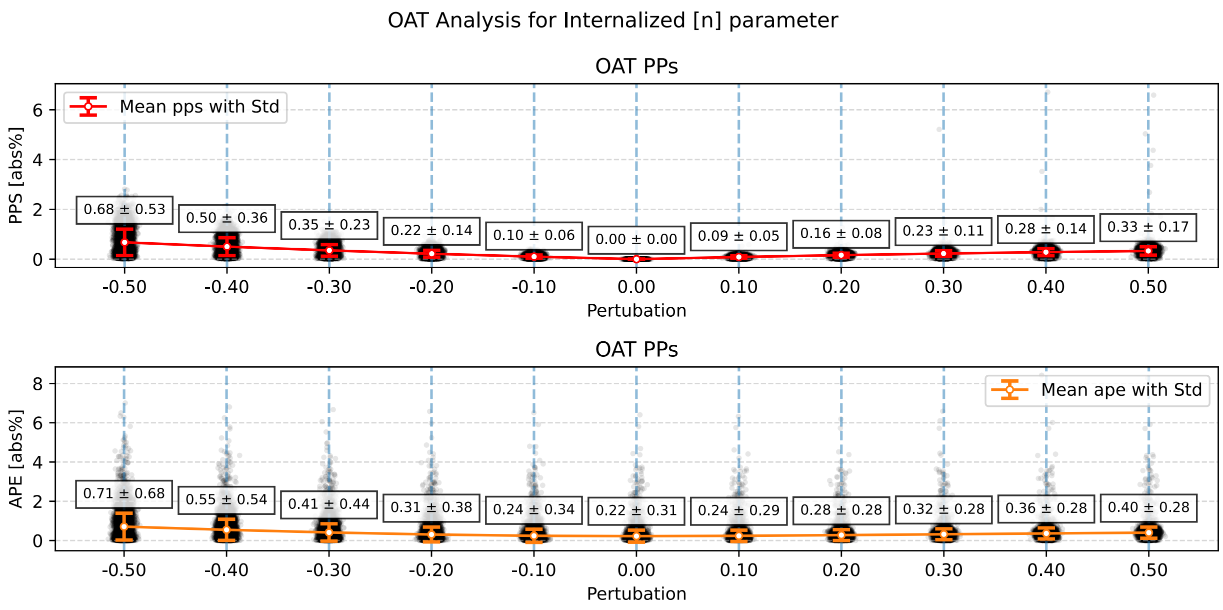


**Supplementary Fig. 3: Prediction dependence on number of internalized LNPs through parameter perturbation.** Top row: shows One at a time (OAT) for the point wise perturbation sensitivity (PPS), strongest dependencies can be seen with negative perturbation reaching 68% prediction change from baseline at negative 50% perturbation. Bottom row: Shows the Absolute Percentage Error (APE) from the simulated test set. The same behavior can be seen here however, reflected in the average relative error, causing a average prediction error up to 72%.

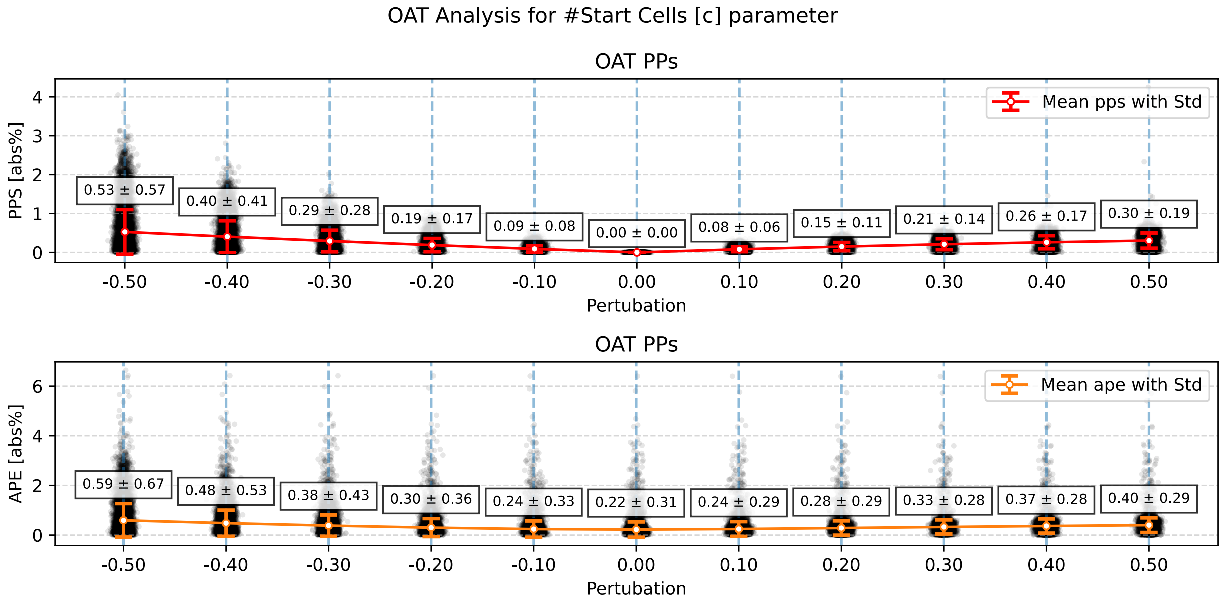


**Supplementary Fig. 4: Prediction dependence on number of Cells through parameter perturbation.** Top row: shows One at a time (OAT) for the point wise perturbation sensitivity (PPS), strongest dependencies can be seen with negative perturbation reaching 54% prediction change from baseline at negative 50% perturbation. Bottom row: Shows the Absolute Percentage Error (APE) from the simulated test set. The same behavior can be seen here however, reflected in the average relative error, causing a average prediction error up to 60%.

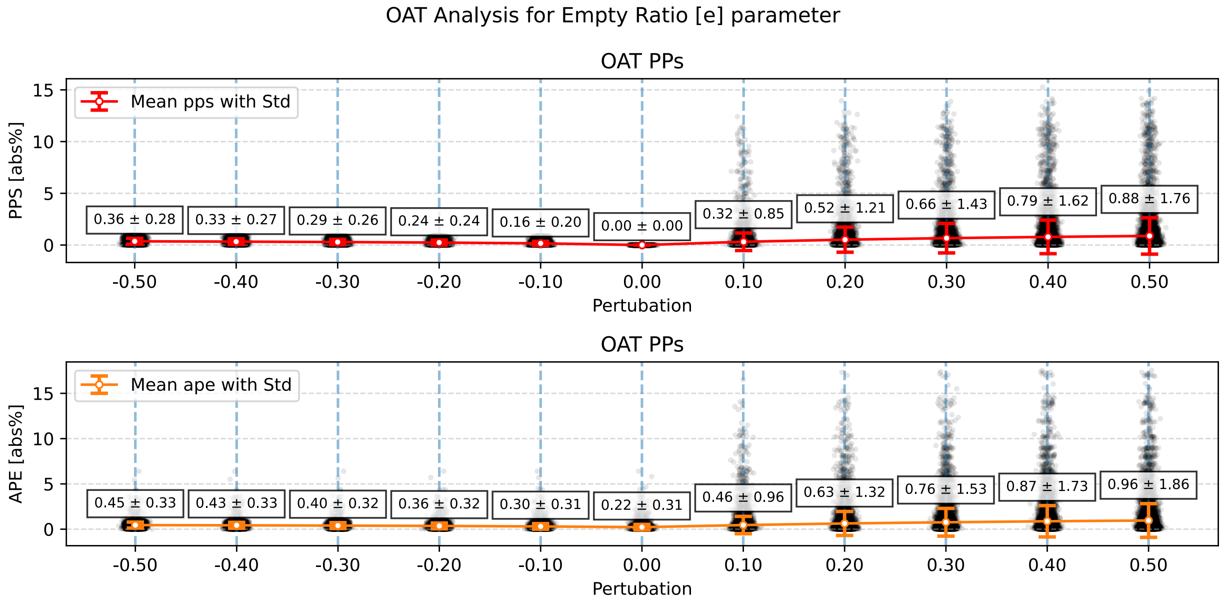


**Supplementary Fig. 5: Prediction dependence on Empty LNPs ratio through parameter perturbation.** Top row: shows One at a time (OAT) for the point wise perturbation sensitivity (PPS), As expected due to the dependencies of Internalized LNPs, wrongly overestimating the empty ratio parameter can have drastic effects on the prediction, with a change up to 90% from baseline at positive 50% perturbation. Bottom row: Shows the Absolute Percentage Error (APE) from the simulated test set. The same behavior can be seen here however, reflected in the average relative error, causing a average prediction error up to 72%.

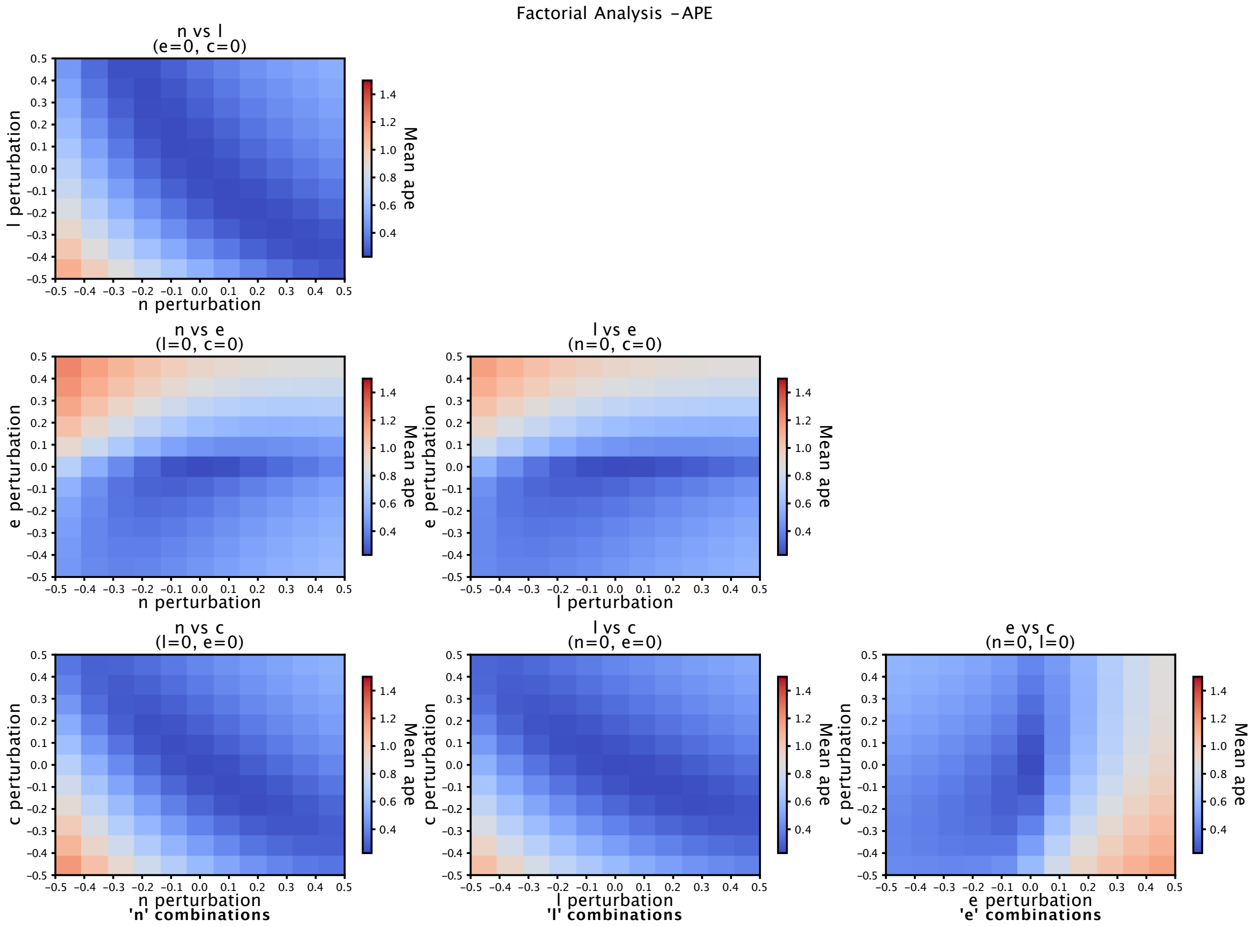


**Supplementary Fig. 6: APE Visualization of the combinatory 2-way perturbation effects across the 4 meta parameters.** Bottom row shows perturbation of *N* Cells on the y axis, with the 3 other parameters at the x axis. In general, the biggest offset Is seen in the edge cases with the negative perturbation expressing the biggest effect. With exception of the Empty LNP ratio which has the opposite effect as seen in SF X. Middle row: depicts the Empty LNP ratio on the y-axis with *N* internalized LNPs and mRNA encapsulations efficiency on the x axis. As seen in the bottom row, the biggest impact is seen in high overestimations of Empty LNP ratios. The top row: Shows the last combination, with mRNA encapsulation on the y-axis and the *N* internalized LNPs on the x axis. The biggest effect here can be seen in the combinatory negative case in the bottom left corner.

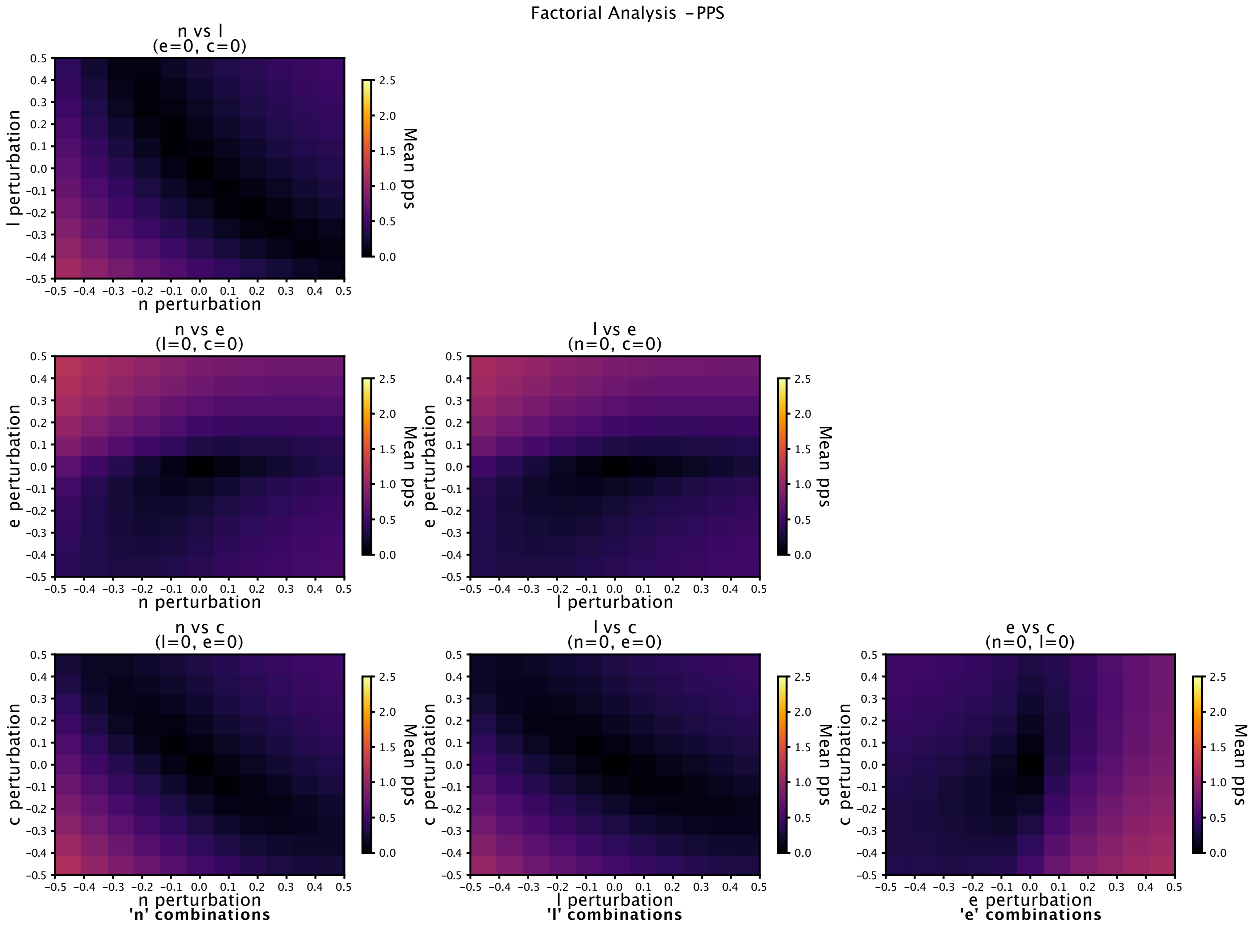


**Supplementary Fig. 7:** Depicts PPS effect from the combinatory 2-way perturbation effects across the 4 parameters. Here the general tendencies follows the same as Supplementary Fig 6.

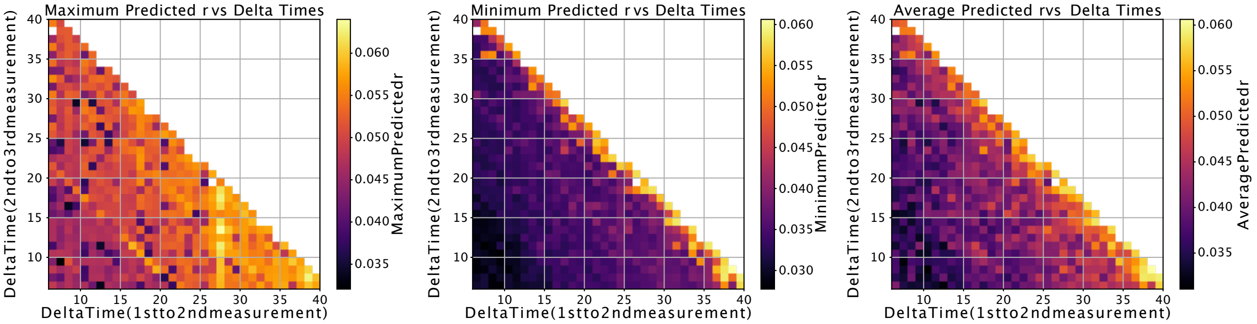


**Supplementary Fig. 8:** Shows the Max**,** Min and Average, predictions of the 3 replicates shown in Figure 3.

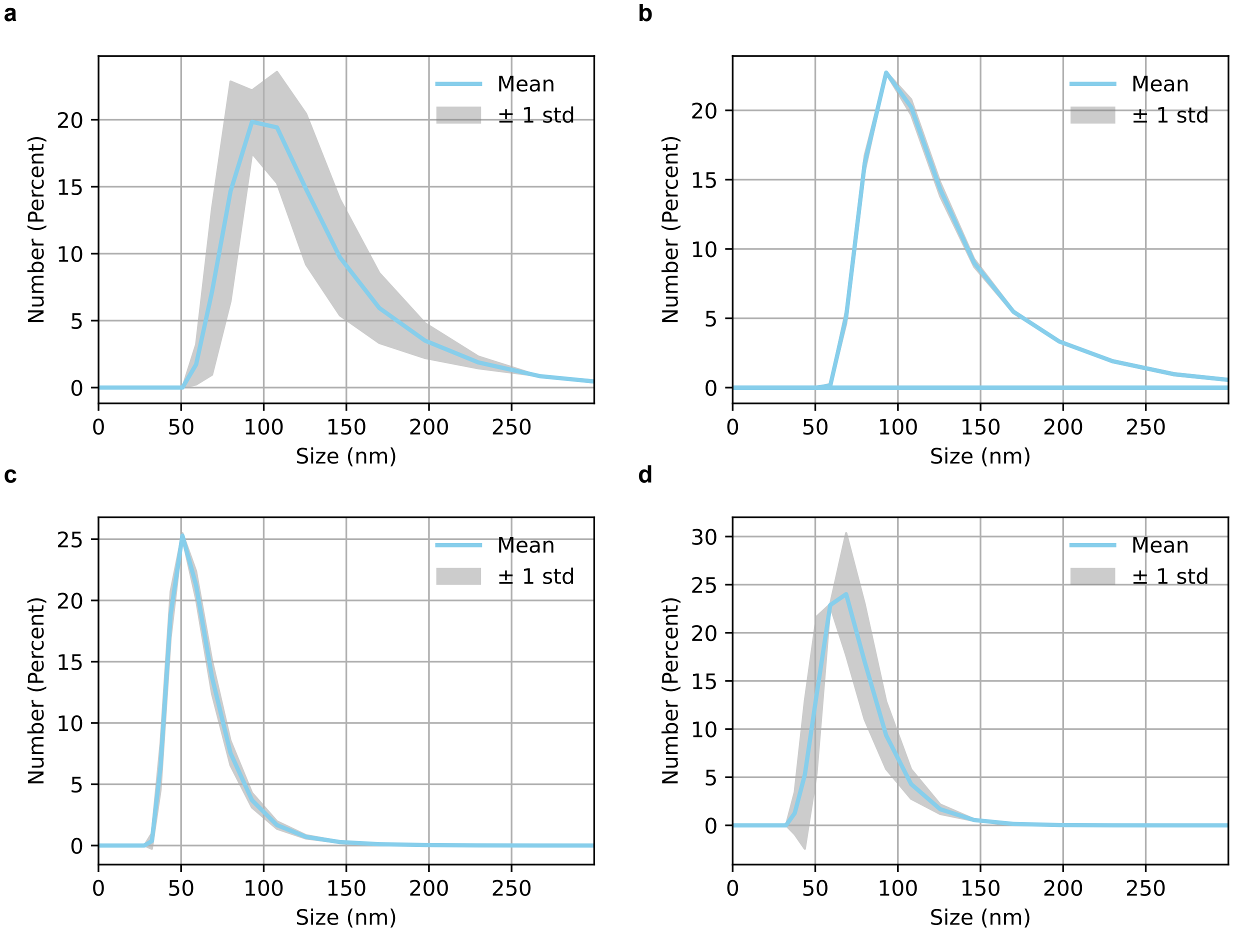


**Supplementary Fig. 9: DLS size characterization of LNPs. Size shown by number. a**, SM-102/cholesterol ATTO655-DOPE_EGFP-mRNA LNP used for internalization studies and EGFP expression studies. **b**, SM-102/cholesterol DSPE-PEG200-biotin ATTO655-DOPE_AF488-EGFP-mRNA LNP used for TIRF single-particle studies. **c**, SM-102/β-sitosterol ATTO655-DOPE_EGFP-mRNA LNP used for internalization studies and EGFP expression studies. **d**, SM-102/β-sitosterol/DSPE-PEG200-biotin ATTO655-DOPE_AF488-EGFP-mRNA LNP used for TIRF single-particle studies.

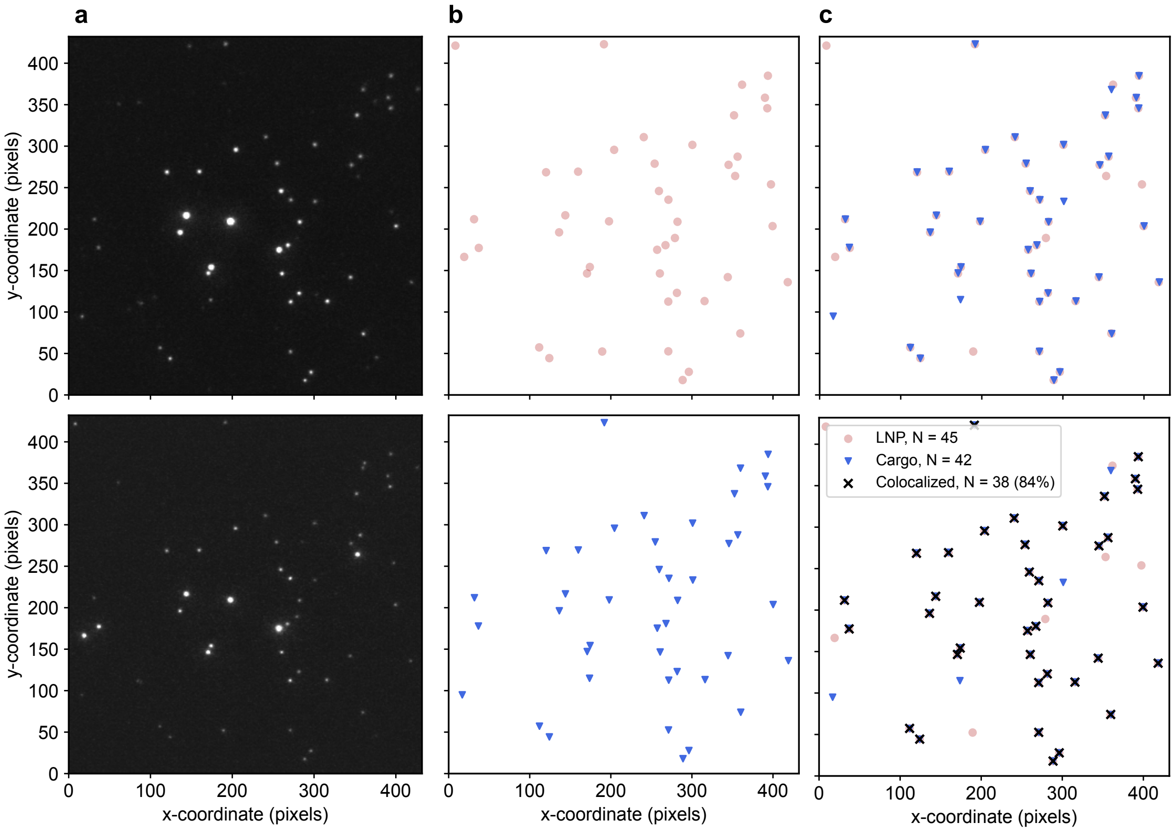


**Supplementary Fig. 10: Quantification of empty LNPs**. **a**, Example images of the membrane channel (top) and cargo channel (bottom). **b**, Example positions of detected particles in membrane (top) and cargo channel (bottom). **c**, Membrane and cargo channel overlaid (top) and where the membrane and cargo channel colocalize (bottom). Across 5 technical replicates and more than $10^5$ LNPs, the percentage of LNPs containing cargo is 86.1 ± 2.0%. *N,* 110635.

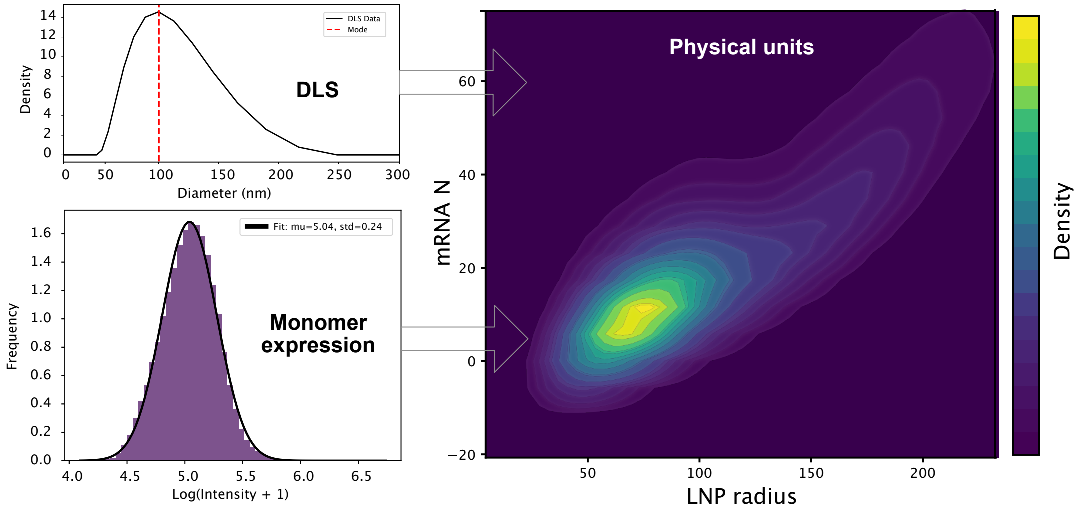


**Supplementary Fig. 11: Estimation of mRNA packing density of single LNPs.** DLS measurements of LNP size are used to convert the measured dual-labelled LNP lipid fluorescence intensity to size. TIRF intensity measurements of mRNA monomer fluorescence intensity are used to convert dual-labelled LNP mRNA fluorescence intensity to mRNA copy number. Each LNP is thus characterized by a size and mRNA count.

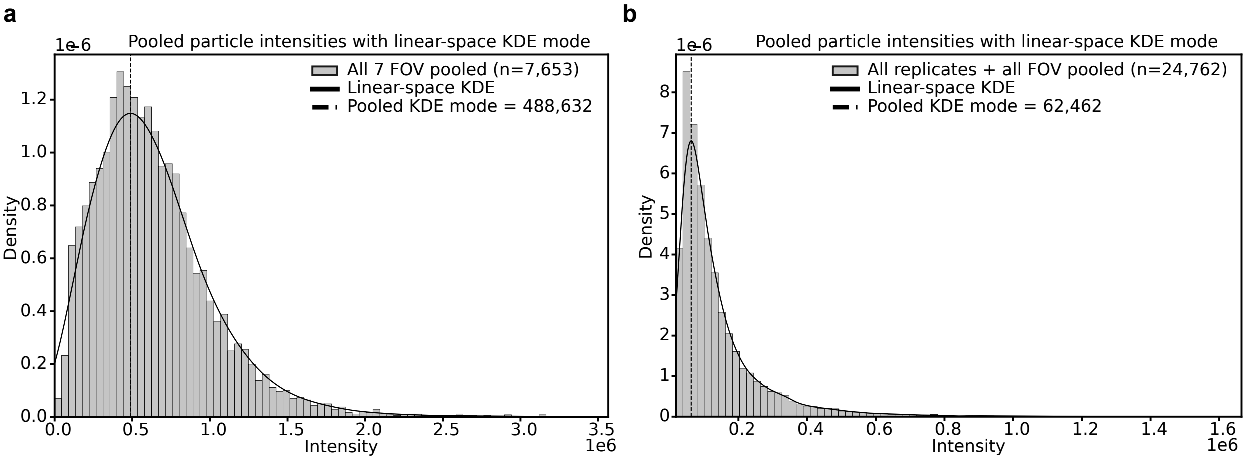


**Supplementary Fig. 12: Quantification of single LNP fluorescence from LLSM data. a,** Intensity distribution is fitted with a KDE to extract the distribution mode for SM-102/cholesterol LNPs. **b**, Intensity distribution is fitted with a KDE to extract the distribution mode for SM-102/β-sitosterol LNPs.

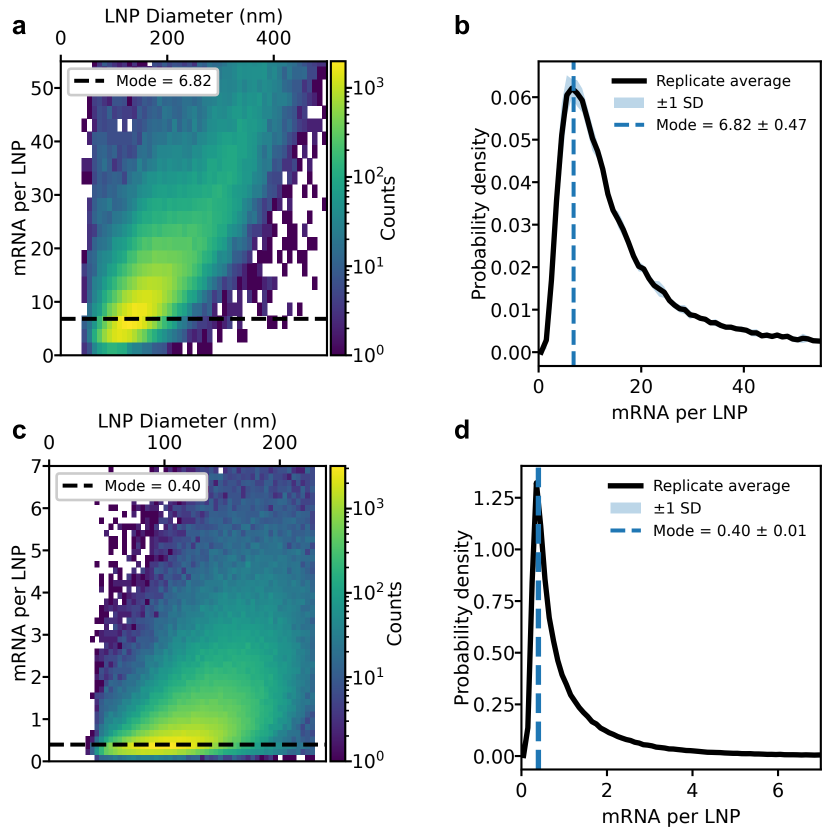


**Supplementary Fig. 13: Distributions of number of mRNA per LNP. a**, Distribuiton of mRNA per LNP versus LNP diameter for SM-102/cholesterol LNP. Data are pooled from four technical replicates. **b**, Distribution of mRNA per LNP for SM-102/cholesterol LNP. A lognormal distribution was fittet to each of four technical replicates to estimate the mode of the distribution. Data are represented as average ± std of four technical replicates. **c**, Distribuiton of mRNA per LNP versus LNP diameter for SM-102/β-sitosterol LNP. Data are pooled from four technical replicates. **d**, Distribution of mRNA per LNP for SM-102/ β-sitosterol LNP. A lognormal distribution was fittet to each of four technical replicates to estimate the mode of the distribution. Data are represented as average ± std of four technical replicates.

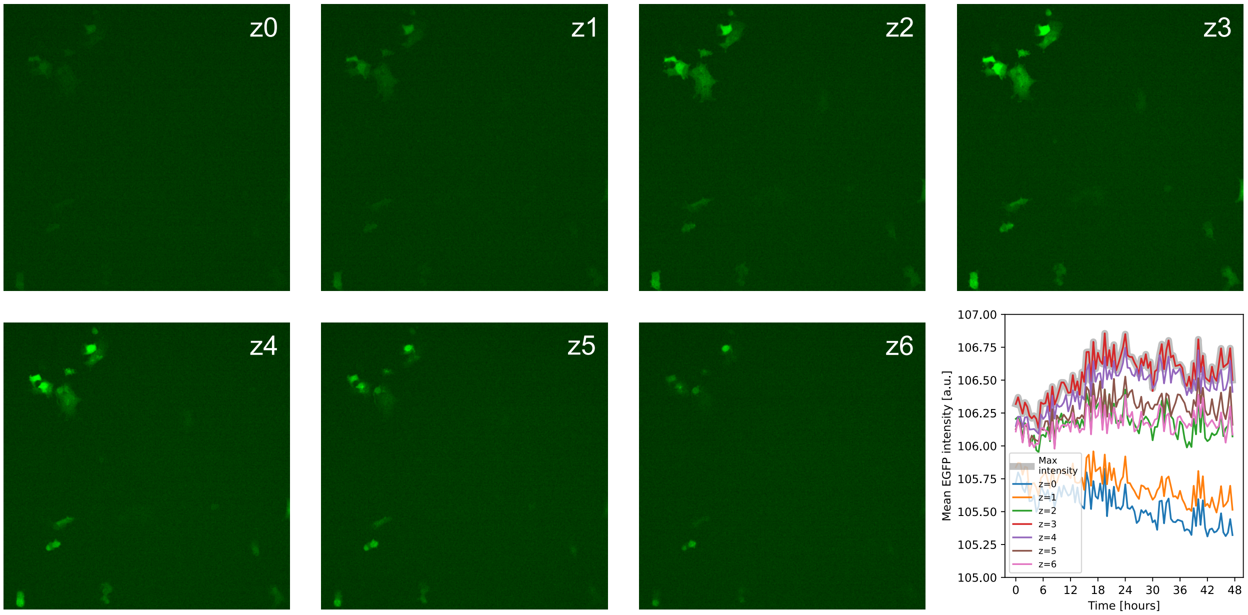


**Supplementary Fig. 14: Selection of max intensity z slice for SDCM data.** All images of last timepoint. Lower right showing evolution over time of mean intensity calculated for the full frame. Mean intensity is highest at closest to the microscopy slide surface where the area of the cell is largest.

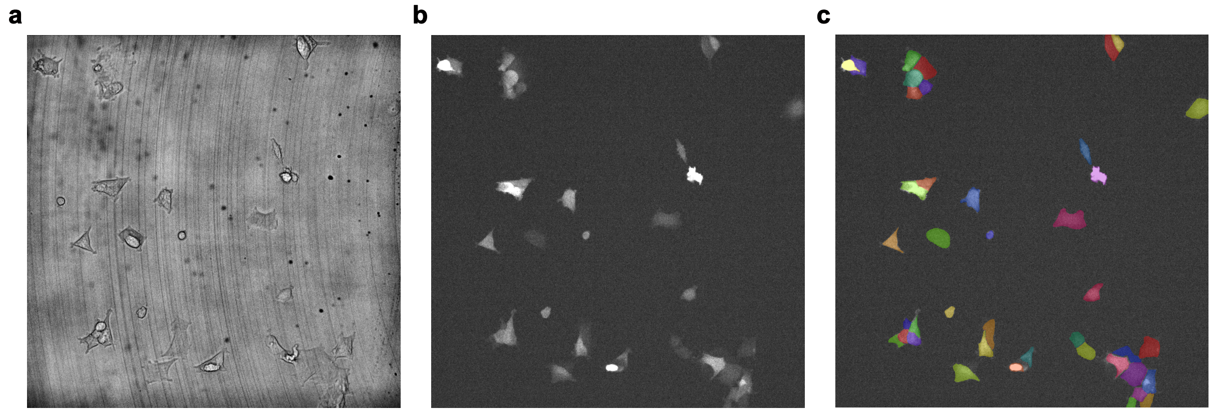


**Supplementary Fig. 15: Segmentation of SDCM data using Cellpose. a**, Brightfield image of HEK293 cells. **b**, Image of EGFP expression in cells in (**a**). **c**, segmentation masks of cells in (**b**) created using Cellpose 3 using EGFP fluorescence signal.

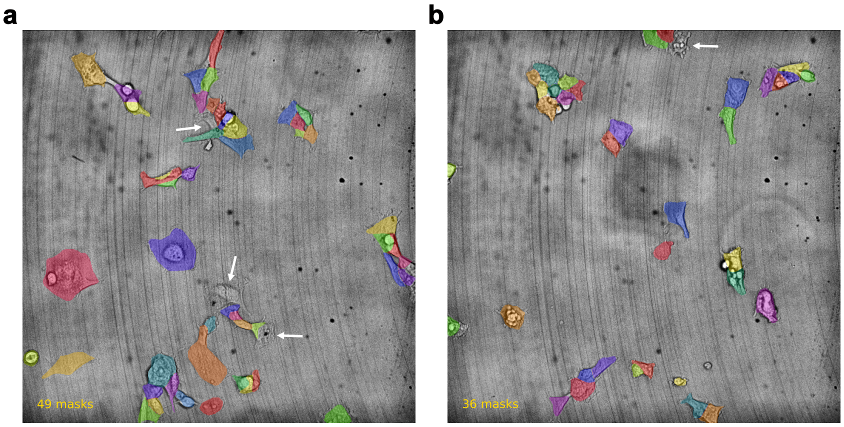


c

| Replicate | 1 | 2 | 3 |
|---|---|---|---|
| EGFP− | 13 | 29 | 34 |
| EGFP+ | 66 | 284 | 418 |
| Total cells | 79 | 313 | 452 |

Negative fraction: 11.1±4.7%

**Supplementary Fig. 16: Counting non-fluorescing cells.** Average percentage negative calculated based on individual replicates. Two exemplary images showing brightfield images of cells and masks of cells created from the EGFP channel (EGFP+). White arrows indicate cells that do not visibly express EGFP (EGFP-).

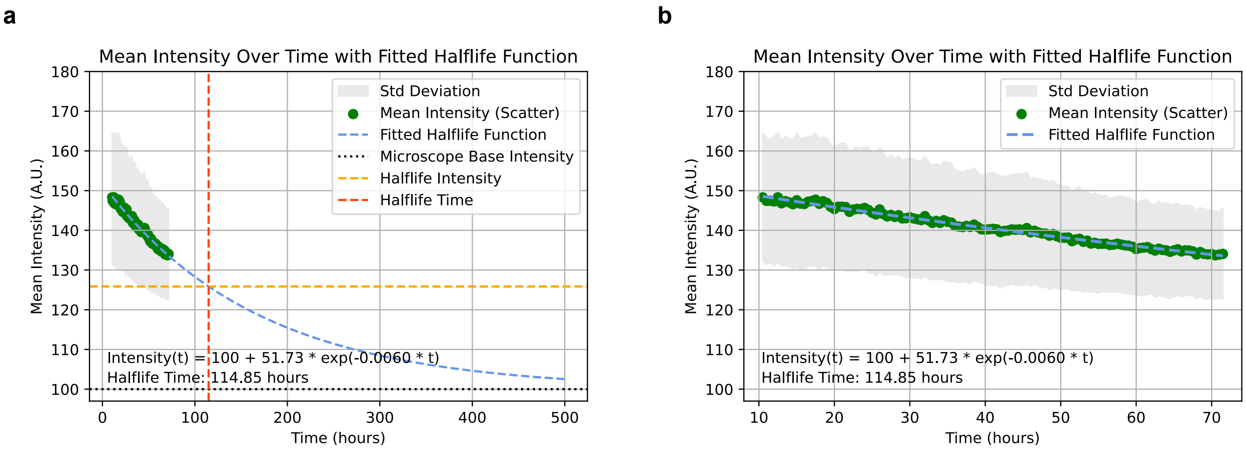


**Supplementary Fig. 17: EGFP degradation rate in live cells**. **a**, Extrapolated fit. **b**, Zoom-in fit on data.

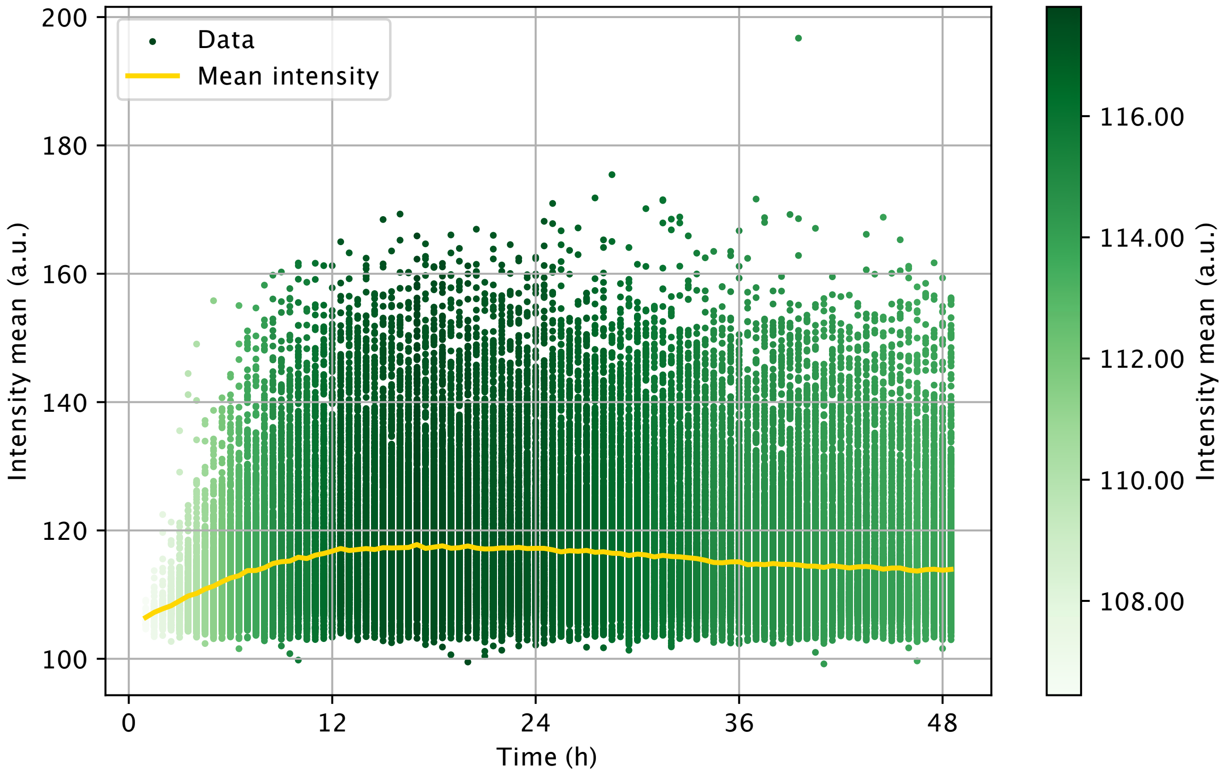


**Supplementary Fig. 18: Single-cell EGFP expression levels from SDCM data.** Each dot represents a single cell expressing EGFP at a single timepoint. *N*=95630.

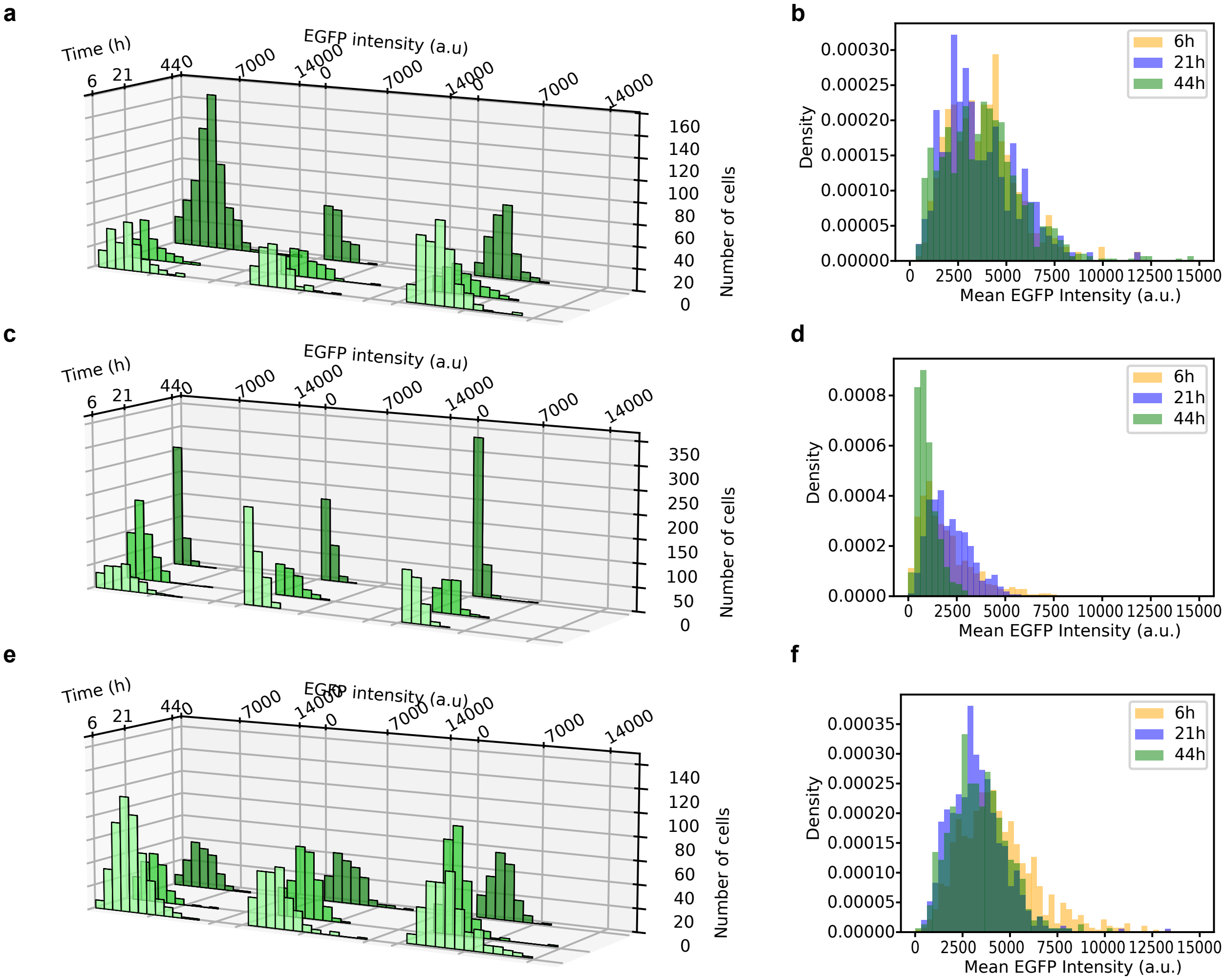


**Supplementary Fig. 19: EGFP intensity distributions from lattice light-sheet experiments. a**, Single-cell EGFP intensity distributions (mean intensity per cell) of HEK293 cells SM-102/cholesterol LNPs. Data from three technical replicates (left to right) at 6 h, 21 h, and 44 h. **b**, Data from **a** with all three replicates pooled for each timepoint. **c,** Single-cell EGFP intensity distributions (mean intensity per cell) of HEK293 cells transfected SM-102/β-siotsterol LNPs. Data from three technical replicates (left to right) at 6 h, 21 h, and 44 h. **d**, Data from **c** with all three replicates pooled for each timepoint. **e**, Single-cell EGFP intensity distributions (mean intensity per cell) of HeLa cells transfected with SM-102/β-siotsterol LNPs. Data from three technical replicates (left to right) at 6 h, 21 h, and 44 h. **f,** Data from **e** with all three replicates pooled for each timepoint.

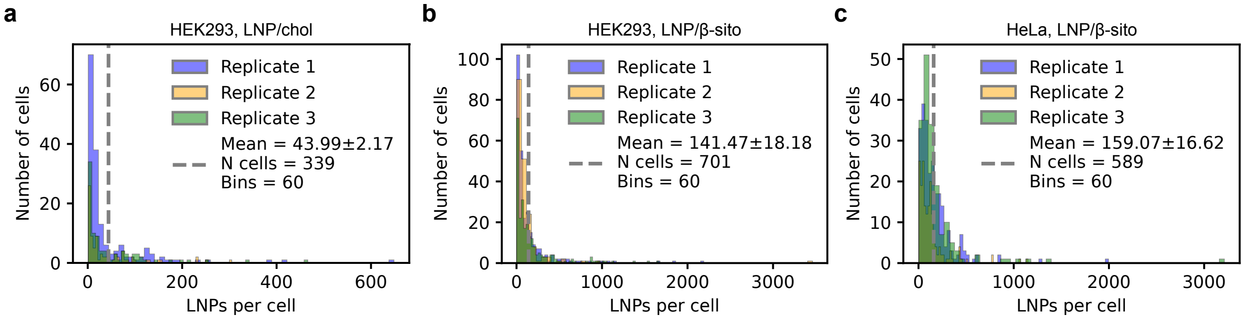


**Supplementary Fig. 20: Quantification of internalized LNPs from volumetric LLSM data normalized to single LNP intensities. a**, Internalization of LNP/chol in HEK293 cells. **b**, Internalization of LNP/β-sito in HEK293 cells. **c**, Internalization of LNP/β in HeLa cells. Mean value depicts the mean ± std of three technical replicates.
